# Classification of protein binding ligands using structural dispersion of binding site atoms from principal axes

**DOI:** 10.1101/2020.12.21.423752

**Authors:** Galkande Iresha Premarathna, Leif Ellingson

## Abstract

Many researchers have studied the relationship between the biological functions of proteins and the structures of both their overall backbones of amino acids and their binding sites. A large amount of the work has focused on summarizing structural features of binding sites as scalar quantities, which can result in a great deal of information loss since the structures are three-dimensional. Additionally, a common way of comparing binding sites is via aligning their atoms, which is a computationally intensive procedure that substantially limits the types of analysis and modeling that can be done. In this work, we develop a novel encoding of binding sites as covariance matrices of the distances of atoms to the principal axes of the structures. This representation is invariant to the chosen coordinate system for the atoms in the binding sites, which removes the need to align the sites to a common coordinate system, is computationally efficient, and permits the development of probability models. These can then be used to both better understand groups of binding sites that bind to the same ligand and perform classification for these ligand groups. We demonstrate the effectiveness of our method through classification studies with two benchmark datasets using nearest mean and polytomous logistic regression classifiers.

## Introduction

Proteins are molecules consisting of chains of amino acids that fold into a 3-dimensional structure that perform biological functions by binding to various chemicals. In protein-ligand binding, the ligand is usually a signal-triggering molecule that binds to a site near the surface on a target protein. A common hypothesis is that proteins which perform similar functions should bind to the same ligands and, as such, have binding sites that have similar shapes. Therefore, analyzing structures of protein-ligand binding sites can be considered as an initial step towards protein function prediction. While studies typically focus initially on binding sites with known binding activity, the common goal is to then utilize these categories of binding sites to predict the binding activity of proteins with unknown function. This problem has many useful applications, such as effective drug discovery with fewer side effects, development of structure-based drug designs, disease diagnosis.

Analyzing these data by hand is time consuming and, as a result, biologists and chemists are tend to work with computer scientists, statisticians, and mathematicians to use bioinformatics techniques to analyze them faster. The Research Collaboratory for Structural Bioinformatics (RCSB) Protein Data Bank (PDB) is the largest data bank that provides information about the 3D structures of proteins and nucleic acid. As of November 10, 2020, there are 170, 597 biological macromolecular structural information files available in PDB and roughly 90% of them are proteins. X-ray crystallography and Nuclear Magnetic Resonance (NMR) are a few common methods used to obtain the protein structure. As of 2003 and 2010, respectively, [1] and [2] showed that as many as 26% of the entries in the PDB have either unknown or putative function. Because much work has been done in this area, those figures continually change due to the discovering of functions and the additions of new structures to the database. An inventory done by [3] shows that there are about 42.53% of PDB entries that were categorized as proteins of unknown functions. By seeing how these figures have changed through past few years, we can understand the amount of research activity that has been going on over these years. As a result, the development of different context-based and structure based method is expanding drastically for prediction of unknown protein function.

Many researchers have conducted ligand-binding protein prediction studies by taking structural information into consideration as an initial step towards protein function prediction. [4] talked about different structure-based approaches used by researchers to predict the binding ligand. Shape based methods, alignment base methods, graph-theoretic approaches, machine learning methods and, model based methods are a few such methods of protein-ligand prediction. In shape based methods, the geometric characteristics being used to determine the similarity of binding sites. Some examples of this type of approach can be found in [5], [6] and [7]. In graph-theoretic methods, they transform protein functions into graphs using different procedures and then use different algorithms to find the relationship between protein structure and the graph. [8] and [9] are some papers that talks about this approach. In machine learning methods, a machine learning environment is adopted for the identification and prediction. [10] and [11] present different machine learning approaches to predict protein-binding ligands. In model-based methods, features of the binding sites for a given group is used to construct a model, and classifications are done based upon the model.

Alignment-based methods provide another popular approach in which binding sites are compared by superimposing them in a pairwise fashion according to some chosen criteria. [12] talked about two web servers and software packages named SiteEngine and Interface-to-Interface (I2I)-SiteEngine for the recognition of the similarity of binding sites and interfaces. [13] talks about a database named SiteBase, which holds information about structural similarity between known ligand-binding sites. For the comparison of these binding sites, geometric hashing was used and the equivalent atom constellations between pairs of binding sites were identified. [14] talks about assessing similarity between pockets in protein binding sites by aligning them in 3D space and comparing the results with a convolution kernel. Then [15] discusses the TIPSA algorithm based on the iterative closest point (ICP) algorithm ( [17]). While many more alignment-based methods exist, these are the key studies that led directly to this current work. While these methods show promising results, they are often computationally expensive to perform and the results can be difficult to analyze since the methods are restricted to utilizing just pairs of binding sites. Using pairwise comparisons of similarity scores rather than structural characteristics of individual binding sites greatly impairs the ability of researchers to develop probability models and machine learning methods for understanding and modeling binding site activity.

On the other hand, when researchers characterize features of solitary binding sites directly, they typically seek to summarize various structural features using scalar quantities. For example, to characterize the size of a binding site, researchers may calculate the volume of the site. While this provides useful information for characterizing the sites, there can be problems with using such an approach. First, for binding sites that are relatively flat, slight changes in the coordinates of even one or two atoms can substantially change a site’s volume, which can, in turn, make the characteristic unstable for use in characterization and modeling. Furthermore, the volume only directly describes information about the size of the region enclosed by the surface of the binding site. Information about the shape and interior structure of the binding site is lost. To combat this type of information loss, researchers can use additional descriptors to quantify these characteristics. For instance, one measure that can describe the shape of a binding site is its sphericity, which is a measure of similarity to a sphere that is proportional to a ratio of a function of the volume of a surface to its surface area. However, this again, fails to directly account for the interior structure of the binding site. A standard measure with which to quantify information about both the size of a binding site and its interior structure is, as used in [15], the radius of gyration, which is the standard deviation of the distances of the atoms to the center of mass of the binding site. Unfortunately, all of these quantities result in a loss of a large portion of information about the structure of the binding sites. While the radius of gyration does at least describe information about the structure of the entire binding site and not just its surface, much information is still lost about the variability within the structure of the binding site because it reduces all of the variability, which occurs in three dimensions, to a single dimension.

These types of univariate measures can certainly be combined together and analyzed either using traditional multivariate analysis methods or various regression analyses to improve our understanding of the structural information of binding sites. However, the relationships between these univariate characteristics are often quite complicated, which can make it more difficult to gain a more complete view of a binding site’s structure.

As such, motivated by the principles of object data analysis (ODA) (See [16]), a more ideal approach is to consider a higher-level representation of a binding site’s structural information that directly incorporates the types of information found in univariate descriptors while also preserving information that can be lost due to condensing such complex information to scalar quantities. In this paper, we will encode the structural information in binding sites as a covariance matrix in a novel way that eliminates the need to align binding sites to place them in a common coordinate system.

The remainder of the paper is organized as follows. In Section 2, we describe the data sets that we will use throughout our study. In Section 3, we present our methodology, including a description of our novel representation of protein binding sites, how we quantify differences between these representations, and a description of the classification procedures we use to evaluate the effectiveness of our representation. In Section 4, we present a detailed analysis of our results, including a discussion about computational costs. Finally, Section 5 contains our conclusions and a discussion of potential areas for future work.

## Data

Motivated by the classification studies of [14] and [15], we decided to focus our attention on two datasets from the literature that consists of a variety of binding sites with varying size, chemical and structural characteristics that all are known to bind to just a handful of ligands so that our eventual classification study could be well-formed and our results could be compared to other methods whose creators had similar goals to ours.

Our first data set, from [18], is known in the literature as the Kahraman dataset. It consists of 100 protein binding sites which bind to one of 10 ligands (AMP, ATP, FAD, FMN, GLC, HEM, NAD, PO4, EST, AND). These ligands vary in size and flexibility. PO4 is the smallest ligand in size and the most rigid molecule. FAD is the largest in size and is the highest in flexibility. Despite this set’s small size, it provides a carefully crafted benchmark set that can be used effectively to demonstrate whether a method can link structure and function via a classification study. The second dataset is called the extended Kahraman dataset in [14]. It consists of 972 protein binding sites, of which the Kahraman dataset is a subset. These sites also bind to one of the above same 10 ligands. Summaries of the data sets are shown in Tables 1 and 2. The relative proportions of each ligand group to the whole data set for the extended data set differ considerably from the original Kahraman set. Most notably, there is a substantially higher proportion of proteins that bind to the ligand PO4 in this dataset.

**Table 1.** Kahraman dataset

|  | AMP | ATP | FAD | FMN | GLC | HEM | NAD | PO4 | STEROID |
| --- | --- | --- | --- | --- | --- | --- | --- | --- | --- |
| Number of Sites | 9 | 14 | 10 | 6 | 5 | 16 | 15 | 20 | 5 |

**Table 2.** Extended Kahraman dataset

|  | AMP | ATP | FAD | FMN | GLC | HEM | NAD | PO4 | STEROID |
| --- | --- | --- | --- | --- | --- | --- | --- | --- | --- |
| Number of Sites | 63 | 78 | 79 | 58 | 88 | 113 | 91 | 389 | 6 |

For the purposes of performing a classification study, one limitation for both data sets is that there are two ligand groups that consist of just a small number of binding sites. In the Kahraman dataset, there are only 2 and 3 binding sites that bind with the ligands AND and EST, respectively. In the extended Kahraman dataset, there are only 2 and 4 binding sites that bind with the ligand AND and EST, respectively. To avoid problems with sample sizes relatively this small and facilitate comparisons with other methods, we condense both ligand groups into one group of steroids, as suggested by [18].

We obtained information about the 3D structures of these proteins from the PDB (Protein Data Bank) that were determined by X-ray crystallography ([19]). To consistently decide what atoms in each protein should be included in binding sites for our study, we adopted the convention of [14], which experimentally determined that all atoms within 5.3 Åof the binding ligand in the crystal structure should be included in a binding site. This definition also facilitates comparisons to [15] and [14]. Unfortunately, though, when we obtained the 3D structure information for the extended Kahraman dataset from PDB, there were 7 binding sites that were removed from the database, resulting in them not being considered in this analysis. While this would prevent us from trying to fully compare our new methodology with other methods, we can still utilize this data to demonstrate the utility of our methods while presenting results for the other methods for reference.

## Methodology

In this research, we approach the ligand-binding protein prediction problem by taking a higher level approach that encodes the structural information found in protein binding sites as a data object in a manner that reduces the amount of information lost compared to using univariate descriptors or structural characteristics. Our approach consists of three main parts: (1) developing a novel representation of binding site structural information as a 3 × 3 covariance matrix that eliminates the need to perform computationally expensive alignment procedures, (2) using properties of covariance matrices to provide a mathematical foundation for quantifying and visualizing dissimilarity between binding sites, and (3) utilizing statistical methods to build nonparametrically defined, empirical probability distributions that both provide insight into the relationship between binding site structure and biological function and allow us to perform classification studies.

### Representation via covariance of distances to principal axes

A natural way to encode information about the structure and size of a point cloud is through the covariance matrix of the coordinates of the points in the cloud. The covariance matrix of the coordinates is

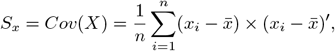

where *x_i_* is a vector containing the 3-dimensional coordinates of the *i*th atom, *n* is the number of atoms in the binding site, 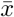 is the 3-dimensional vector of the coordinates for center of mass of the atoms in the binding site, and ′ denotes the matrix transpose. For a 3-dimensional point cloud, *S_x_* is a 3 × 3 symmetric positive definite (SPD) matrix. Since the set of all of these matrices is a metric space, we could utilize a distance defined on this space to quantify the degree of dissimilarity between two point clouds, regardless of the number of points present in each cloud. In this application, the points are each atom in a binding site. Unfortunately, we immediately run into a critical problem with using this as a representation of binding site structure. The coordinates for the locations of the atoms are provided with respect to their locations in the crystals of the full proteins using arbitrary vector bases, so no two binding sites share the same *x, y*, and *z* axes. As a result, the covariance matrices characterize completely different directions of variability, which prevents us from using the distance between two covariance matrices as a useful measure of dissimilarity. Because of this, it would initially seem that we would need to appeal to alignment-based methods to obtain common coordinate systems.

However, motivated by the *PrincAxis* similarity measure used by [14], which quantifies the differences in the lengths of the principal axes of two binding sites, we turned to principal component analysis to identify the principal axes, which are the three orthogonal directions passing through 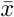 along which variability in the atom coordinates is maximized. An example showing the principal axes for a binding site is shown in Fig 1. The axes are obtained by finding the eigenvectors corresponding the the eigenvalues, in decreasing order, of *S_x_*.

**Fig 1.**
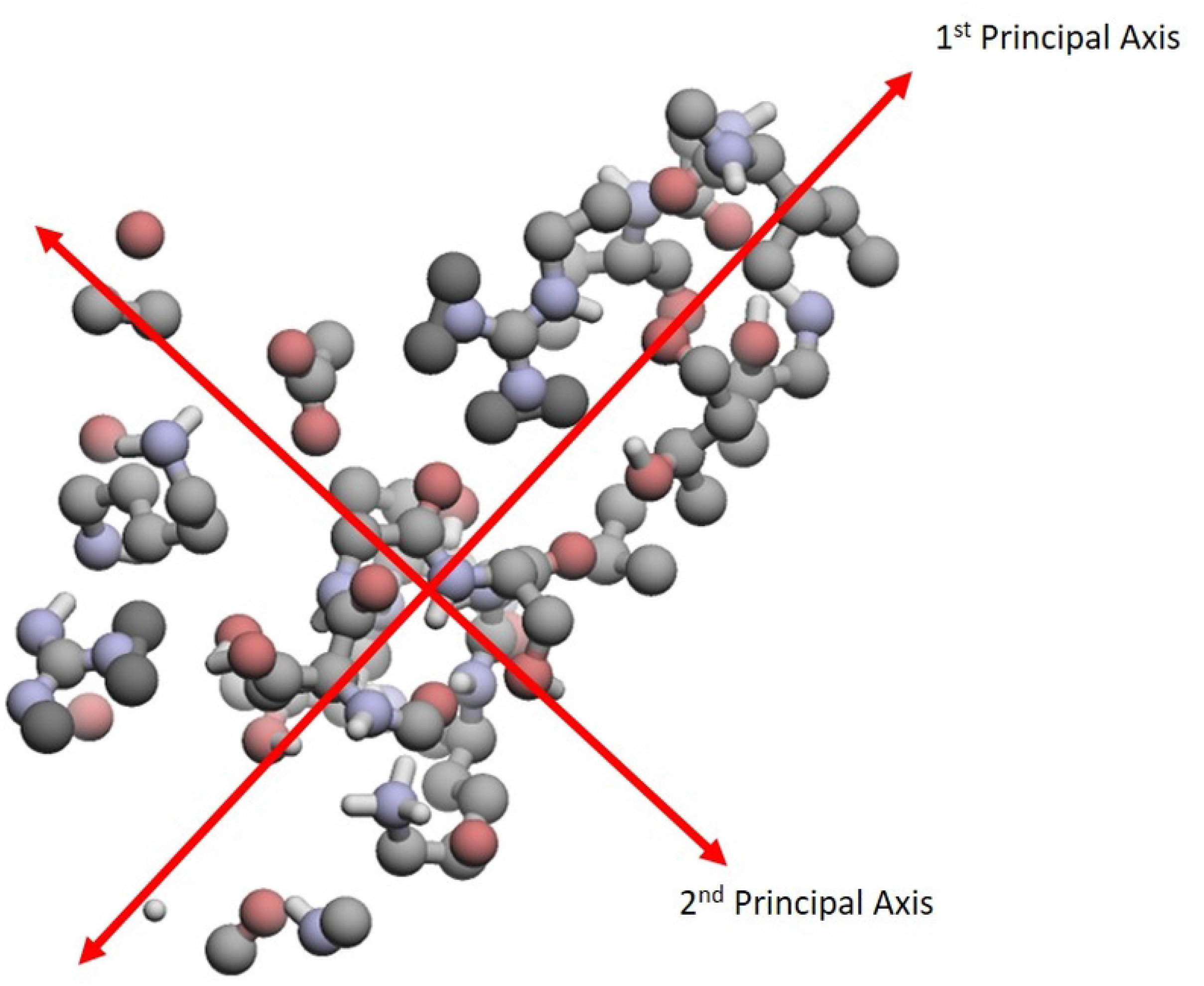
Orthogonal principal axes of 1AYL-ATP. This is an image of the binding site 1AYL-ATP. It illustrates where each principal axis is located. Note that the third principal axis is orthogonal to the page.

Since all binding sites have three principal axes, they provide a common coordinate system for encoding the structural variability that we can subsequently use to compare binding sites. To do this, we focus on the distances of each atom to the three principal axes, where the distance of an atom to a principal axis is defined to be the Euclidean distance between the atom’s coordinates and its projection onto the principal axis. Once we have the distances for all atoms, we can then construct the covariance matrix of these distances, which we use as our data object for encoding the structural information from binding sites. For convenience, we will, from here on out, refer to this representation as the Covariance of Distances to Principal Axes (CDPA).

We will now describe the details of this process mathematically. Let *d_kj_* denote the distance of the *k^th^* atom to the *j^th^* principal axis, where *k* = 1, 2,…, *n_i_, j* = 1, 2, 3, and *n_i_* is the number of atoms for the *i*^th^ binding site, for *i* = 1, 2,…, 100 or *i* = 1, 2,…, 972, depending on the data set we use. Then, the distance matrix *d_i_* for the *i*th binding site can be represented as

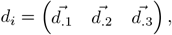

where 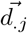 is the vector of the distances of all *n_i_* atoms to the *j^th^* principal axis. Our final data objects are then defined as

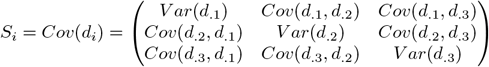

for *i*=1, 2,…, 100 or *i*=1, 2,…, 972.

As an illustrative example, we consider the PO4 binding site of the protein 1cbq, which is found in the extended Kahraman dataset and consists of only 8 atoms. The distances of each atom to all three principal axes for this binding site are shown in matrix form as

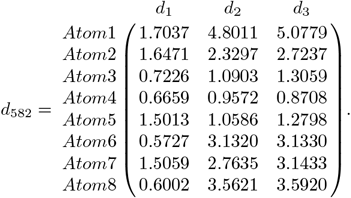

Then, the covariance matrix for ligand-binding site, 1cbq-PO4 can be shown as

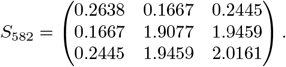

### Quantifying differences between binding sites

In order to classify binding sites according to their binding ligand using CDPA, we need to be able to quantify differences between binding sites. To do so, we need to utilize properties of the space of 3 by 3 SPD matrices. While this space, itself, is not a vector space, it is a submanifold of the space of 3 by 3 symmetric matrices, which is a Euclidean vector space. As such, it inherits the Euclidean distance

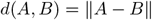

for symmetric matrices *A* and *B*, where 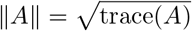. However, since the Euclidean distance alone fails to take the covariance structure of the data into account, we must instead use the Mahalanobis distance between two matrices *A* and *B* to get a more meaningful measure of dissimilarity between binding sites. In order to calculate the Mahalanobis distances, means and covariances of *S_i_* are required, but since each *S_i_* is already a matrix, we must first vectorize the observations. To do so, we utilize the vectorized form vecd(*A*) described in [20] and [21], which is calculated as

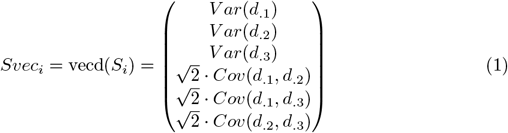

The last three entries of Eq (1) are multiplied by 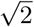 so that the Euclidean distance between any two observations remains the same whether in matrix or vector form. In other words, the Frobenius norm of the matrix will be equal to the norm of the vectorized form of the matrix. That is, trace(*S_i_*) = (*Svec_i_*)′(*Svec_i_*).

With this form, we can obtain the mean vector and covariance matrix for each group of binding ligands in the standard way to obtain 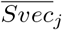 and 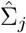, respectively, for *j* = 1,…, 9. This allows us to calculate the Mahalanobis distance from each binding site i to the mean of ligand group *j* as

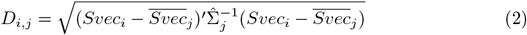

where, *i*=1, 2,…, 100 or *i*=1, 2,…, 972 and *j*=1, 2,…, 9.

Using these calculated Mahalanobis distances, we can directly examine the relationships between the binding sites and the means of each group by comparing the distributions of the Mahalanobis distances for binding sites in group *j* to the mean of group *j* with the Mahalanobis distances for binding sites in other groups to the mean of group *j*. We show an example of histograms for these distributions in Fig 2 and 3 for the PO4 group. It is clear that nearly all of the PO4 binding sites are quite close to the mean of PO4 while the binding sites from other groups generally are further away from the mean of PO4. This, along with the similar results from the other groups, suggests that we can use these distributions as nonparametric models for each group as a a basis for performing classification.

**Fig 2.**
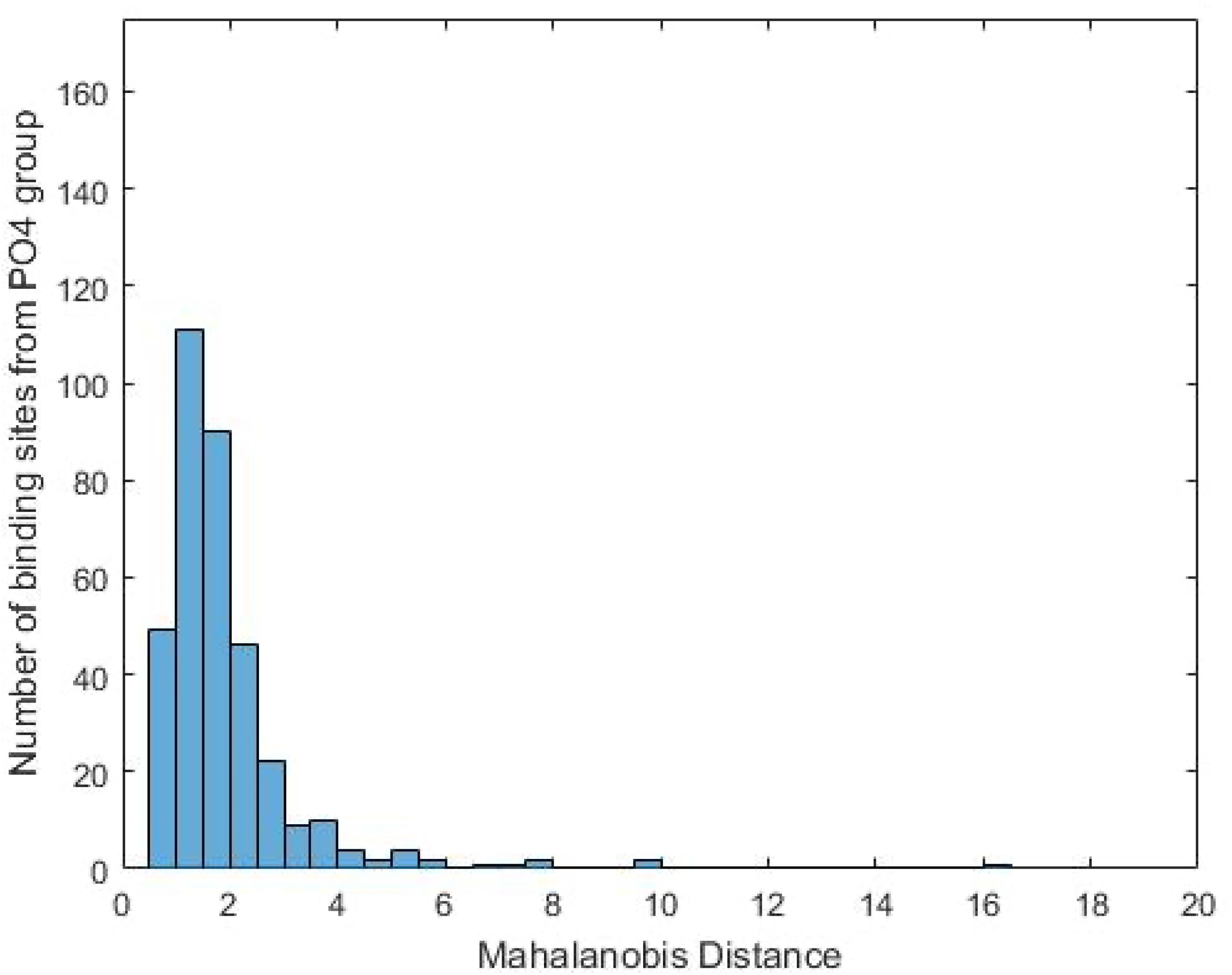
Mahalanobis distance from each binding site in the PO4 group to the mean of the PO4 group. This histogram shows an example of a distribution of the Mahalanobis distances for binding sites from a ligand-binding group to the mean of the same group.

**Fig 3.**
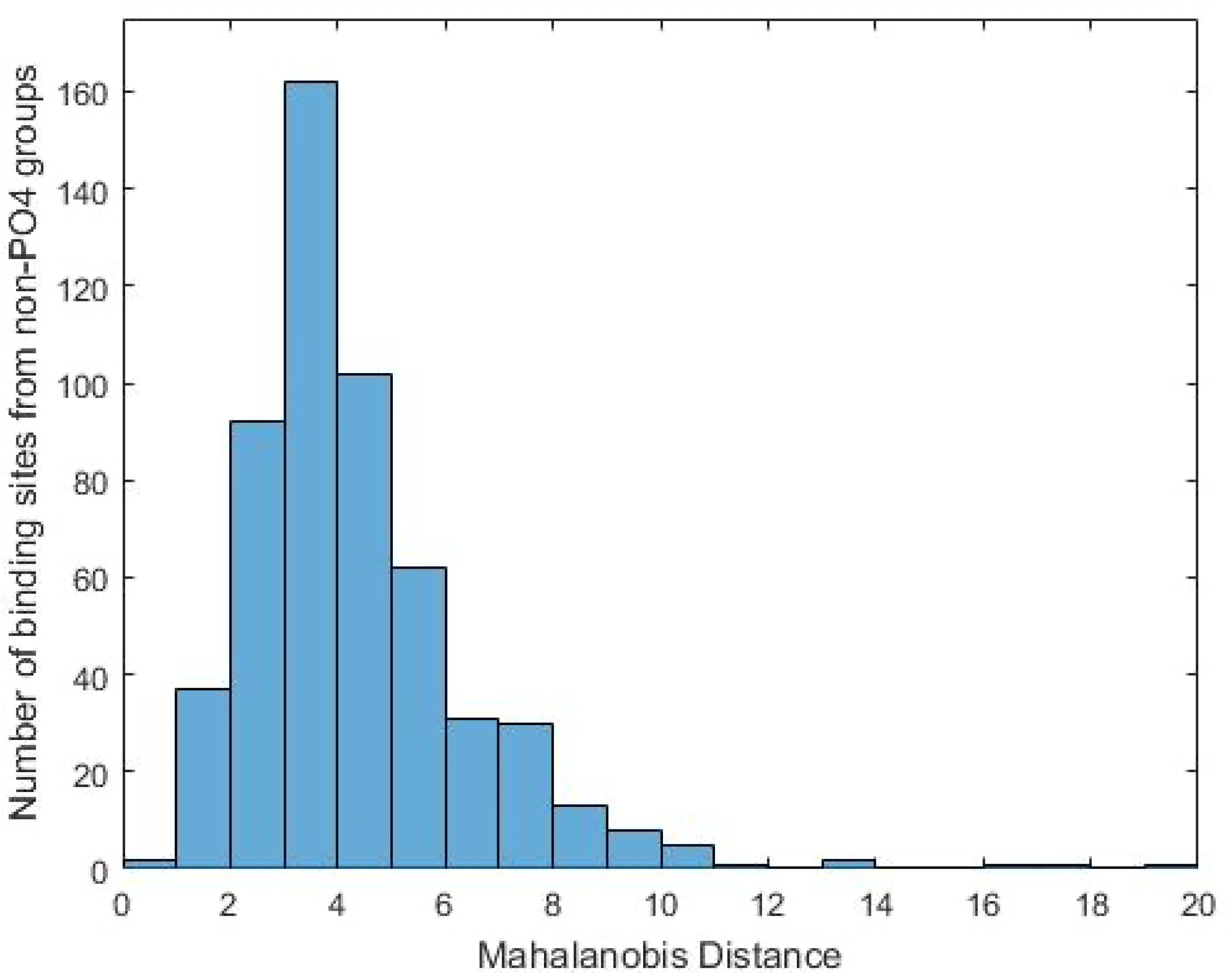
Mahalanobis distance from each binding site in the non-PO4 groups to the mean of the PO4 group. This histogram shows an example of a distribution of the Mahalanobis distances for binding sites in other groups to the mean of the current group.

### Classification and validation methods

We use two procedures to classify the binding ligand of each binding site. The first is the nearest mean classifier. It is a form of nearest neighbor classification, which was utilized by [15] and [14]. However, while the latter method requires all observations to be compared to each other in a pairwise fashion, the nearest mean classifier requires only for each observation be compared to the mean of each group. For a fixed observation *i*, we compute *D_i,j_*, as calculated in Eq (2), for *j* = 1,…, 9 and assign it to group *j*′ if

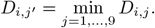

Unlike nearest neighbor classification, this method allows us to explicitly utilize the mean and covariance structure of each binding site and is based upon the models like those shown in Fig 2 and 3. An advantage of this classifier is that it is simple, so we can nearly directly determine the utility of CDPA and the Mahalanobis distance models with minimal impact from the classification scheme.

The second method we use is polytomous/multinomial logistic regression, which allows us to model the probability that binding site *i* binds to ligand group *j*′. As predictor variables, we use the *D_i,j_* so that the probability depends not just on what group is closest to observation *i*, but how close observation is to every group. This allows us to use both the variability within groups and between groups to classify each binding site. If we denote the predicted probability that observation i belongs to group *j* as *P_i,j_*, then we assign observation *i* to group *j*′ if

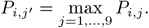

An additional advantage of logistic regression is that these predicted probabilities also give us a measure of certainty regarding the classifications. If *P_i,j′_* is high, then we should feel more certain that the classification is correct, whereas *P_i,j′_* taking a low value means that the logistic regression model has trouble distinguishing group *j*′ from the others for the observation.

To validate the classification scheme, we initially considered the classical leave-one-out cross validation scheme, but, unfortunately, many of the ligand groups in both data sets contain very few binding sites, so leaving out even one observation destroys covariance structure of the group, rendering the entire analysis unstable. As an example, we visualize the observations in the steroid group in 3 dimensions using multidimensional scaling (MDS) in Fig 4.

**Fig 4.**
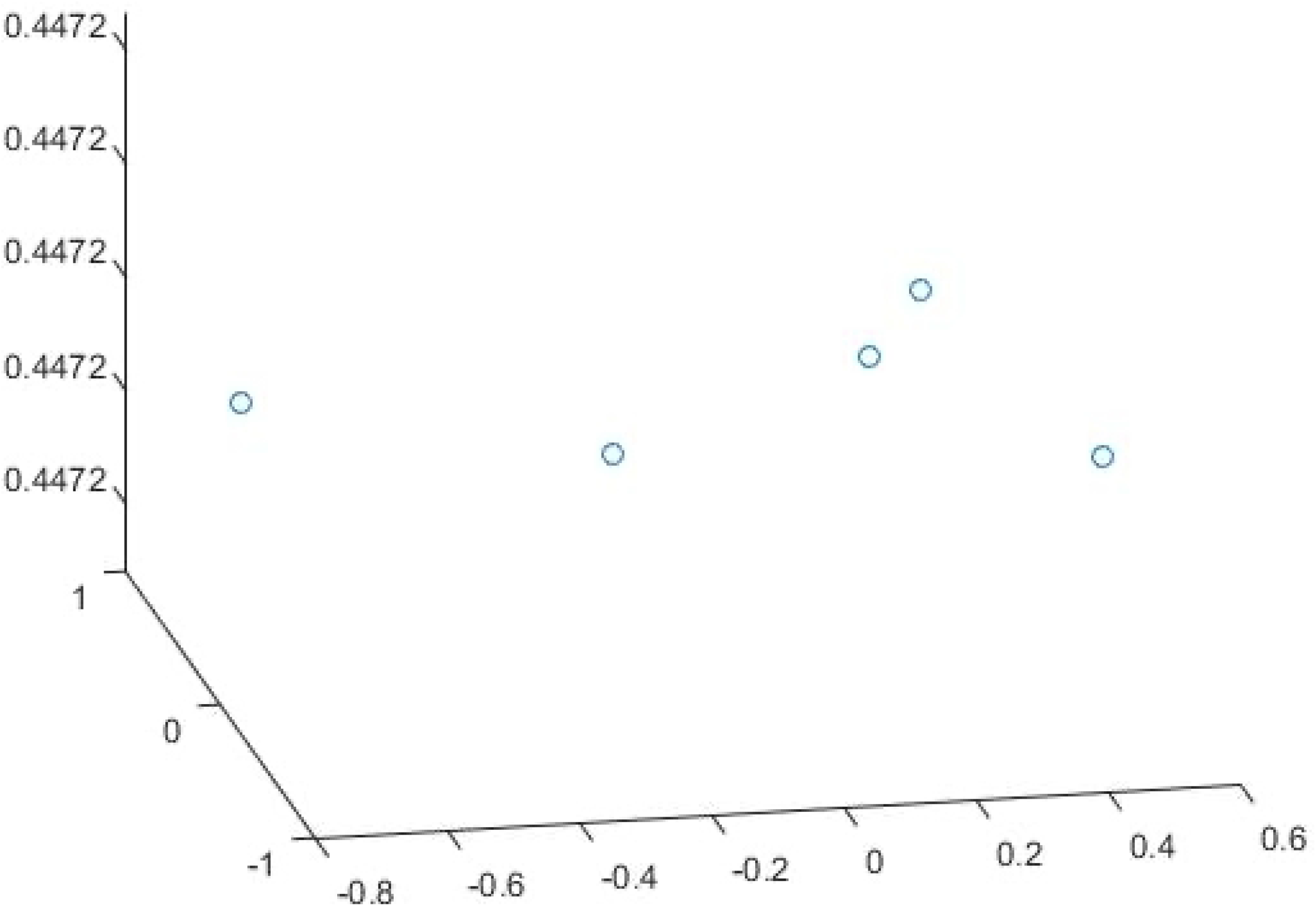
MDS plot in 3 dimensions for the Steroid group in Kahraman dataset. If even a single observation is removed, the mean and covariance structure for the entire group will be radically altered, which makes leave-one-out cross validation fail.

Instead, we can simulate a validation dataset, allowing us to use the entire original dataset as a training set, leaving the means and covariances unaltered. To do this, we simulate testing data by adding noise to each atom coordinate. First, the initial data are read-in, and the atom coordinates for given binding sites are measured. For instance, suppose the coordinates of atom *j* are given as 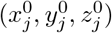. Then, a small amount of 3 dimensional Gaussian noise (*τ*) is added to the coordinates of each atom to perturb the data and can be represented as

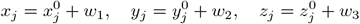

where *w*_1_, *w*_2_, and *w*_3_ are independent *N*(0, *τ*) random variables. CDPA and classification is then repeated using the noise-altered data with respect to the models formed using the original training set. For a given value of *τ*, we repeat this procedure 100 times so that we can detect the influence of individual observations versus a particular realization of the noise. We repeat this procedure over a fine grid of values for *τ* from just above 0 to 1. Given that all atoms in each site are within 5.3 Åof the binding ligand, values of *τ* near the upper end of this range may significantly alter the structure of the atoms in the binding site.

## Results

We now present the results of performing classification studies for both datasets using CDPA and discuss the particular challenges involved in working with each dataset.

### Ligand classification for Kahraman dataset

We first consider the smaller Kahraman dataset so that we can compare our results to those of both TIPSA and sup-CK. First, we visualize the entire data set in 3 dimensions using MDS in Fig 5. The plot shows a clear separation between most of the groups and we can see that the covariance structure for the groups differ from each other considerably. This further supports why it is imperative to use Mahalanobis distances rather than just Euclidean distances between the binding sites when analyzing the data.

**Fig 5.**
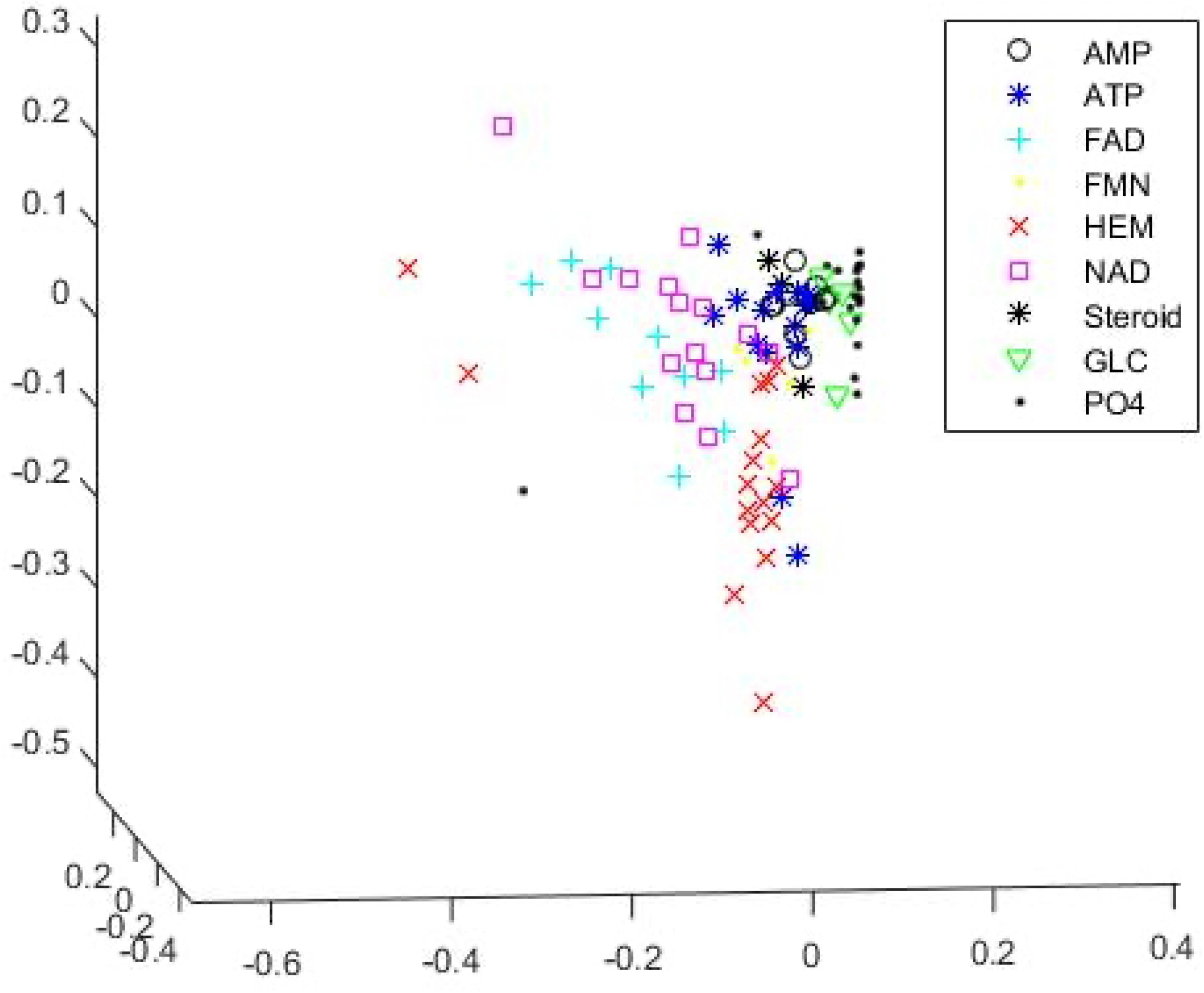
MDS plot in 3 dimensions for the Kahraman dataset.

A summary of classification results is shown in Table 3. The classification errors are the proportions of binding sites that were classified to the incorrect ligand group for each method used. Note that we only report the results for nearest mean classification for CDPA for this dataset in the table. This is because, since we must use one ligand group as a baseline category, we have 8 logistic models, each of which contains 8 parameters, which gives us a total of 80 parameters that must be estimated. This leaves us with just 20 degrees of freedom out of the 100 observations. As a result, even though the fitted model produced a classification error of 0%, this is speaks only to drastic overfitting rather than to any benefit of the methodology.

**Table 3.** Results for nearest mean classification for the Kahraman (5.3 Å) dataset.

| Method | Classification Error |
| --- | --- |
| TIPSA:TI | 0.43 |
| Gyr | 0.54 |
| TIPSA:TI+Gyr | 0.29 |
| $Sup - CK_L$ | 0.27 |
| **CDPA with nearest mean classification** | **0.15** |

In the table, we see that CDPA with the nearest mean classifier has roughly half the classification error as the leading versions of both TIPSA and sup-CK. While this indicates that CDPA outperforms the leading methods for this dataset, the most important takeaway from these results is that it is clear that the CDPA representation is able to effectively encode useful information about the structures of the binding sites. We also include results for a few variations from the TIPSA study to further highlight the importance of taking the covariance structure of the binding sites into account. TIPSA:TI uses the Tanimoto Index (also known as the Jaccard Score), which references only the proportion of atoms common to both binding sites in a pair, as a similarity measure. Gyr is the difference between the radii of gyration for two binding sites, so it utilizes a one dimensional measure of the variability of atoms within binding sites as a dissimilarity measure. While TIPSA:TI performs better than Gyr by themselves, Ellingson and Zhang (2013) showed that using Gyr in combination with the Tanimoto Index improved the results considerably. This provides further evidence that it is important to utilize information about the covariance structure of the binding sites.

A confusion matrix showing a detailed breakdown of the results for CDPA is shown in Table 4. From this, we can see that 9 of the 15 misclassified binding sites are assigned to the HEM group. This makes sense based on Fig 5, since we see that, while most of the variation in the HEM group is along the vertical axis, there are two binding sites in the group that are outliers, though still within the scope of the variation exhibited by the entire dataset, that significantly affect the covariance matrix for HEM. This thusly results in some binding sites from other groups being misclassified as HEM sites.

**Table 4.** Correctly classified and misclassified number of binding sites in each ligand group for Kahraman dataset using nearest mean classification

|  | AMP | ATP | FAD | FMN | HEM | NAD | Steroid | GLC | PO4 |
| --- | --- | --- | --- | --- | --- | --- | --- | --- | --- |
| AMP(9) | **8** | 1 | 0 | 0 | 0 | 0 | 0 | 0 | 0 |
| ATP(14) | 1 | **10** | 0 | 0 | 3 | 0 | 0 | 0 | 0 |
| FAD(10) | 0 | 0 | **7** | 0 | 3 | 0 | 0 | 0 | 0 |
| FMN(6) | 0 | 1 | 0 | **4** | 0 | 0 | 0 | 0 | 1 |
| HEM(16) | 0 | 0 | 0 | 0 | **16** | 0 | 0 | 0 | 0 |
| NAD(15) | 0 | 0 | 1 | 0 | 2 | **12** | 0 | 0 | 0 |
| Steroid(5) | 0 | 0 | 0 | 0 | 0 | 0 | **5** | 0 | 0 |
| GLC(5) | 0 | 0 | 0 | 0 | 0 | 0 | 0 | **5** | 0 |
| PO4(20) | 0 | 1 | 0 | 0 | 1 | 0 | 0 | 0 | **18** |

To perform model validation, we added noise to the data, as described in the previous section. A plot of the classification error as a function of noise level *τ* is shown in Fig 6. The dark line represents the average classification error over the 100 replications while the shaded region surrounding it provides 95% confidence bands. As expected, as *τ* increases, the performance of the method degrades. However, we can see that the classification error is still better than or comparable to the other leading methods for the Kahraman set (while they use the original data) for values of *τ* up to roughly 0.3. Additionally, the confidence bands are thinner for values of *τ* less than 0.3, and only become more consistently thick for higher levels of noise, which appear to have significantly altered the structures of the binding sites. Even for those largest values, though, CDPA with the nearest mean classifier still performs comparably to TIPSA:TI.

**Fig 6.**
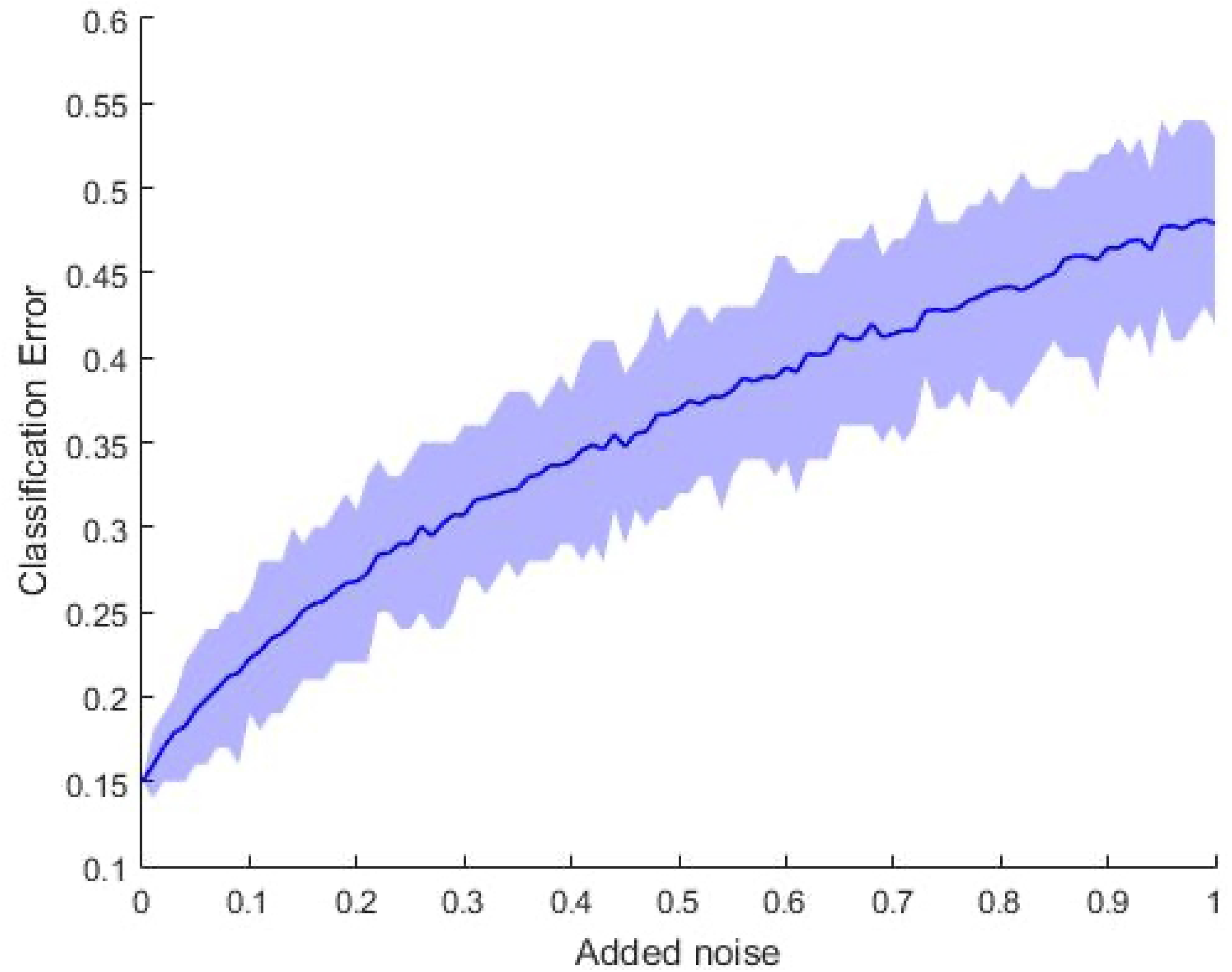
Classification error with respect to different noise levels for the Kahraman dataset

### Ligand classification for extended Kahraman dataset

The analysis proved to be more complicated for the extended Kahraman dataset. Since 7 out of 972 binding sites from the extended Kahraman dataset were unable to extract from the PDB, only 965 observations were available for analyzing the data and performing classification. We visualize the full data set using MDS in 3 dimensions in Fig 7. Since it shows that most of the binding sites are concentrated closely together, and only some of the binding sites are scattered over a large region (most of which are in the PO4 group), it is clear that the data needs to be cleaned in order to be properly analyzed. Indeed, when we performed an initial classification study using the nearest mean classifier, we saw a classification error of 0.5389.

**Fig 7.**
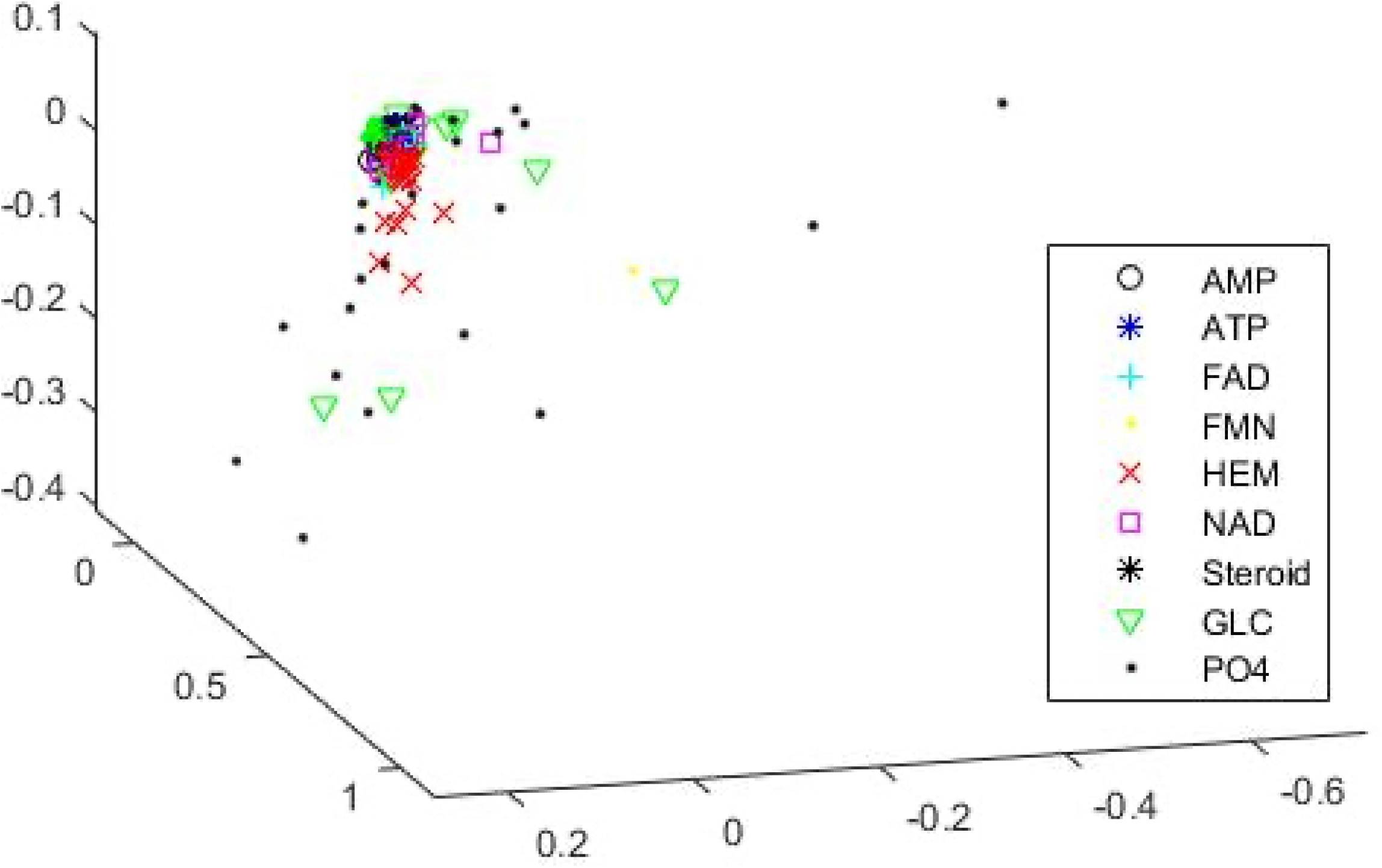
MDS plot in 3 dimensions for all ligand groups of the extended Kahraman dataset before cleaning the data.

We looked more closely at the data to determine why the method produced such a high error rate and calculated the covariance matrix 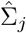 for each group. We found that the FMN and PO4 groups have extremely high levels of variation in at least some variables. Their covariances are shown below.

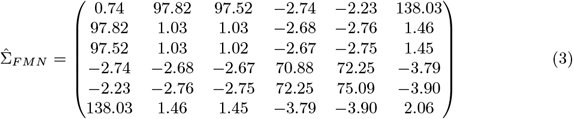

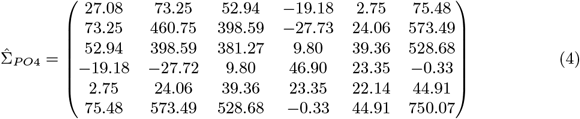

There are higher variations present in these ligand groups. Therefore, to check which binding sites affect the covariance structure by behaving differently in each of these groups, we constructed MDS plots in 3 dimensions for both, which are shown in Fig 8 and 9. In the FMN group, it is clear that one observation differs greatly from the others, which is what causes the high amounts of variability for two of the variables in Eq (3). Likewise, matrix Eq (4) for the PO4 group shows that a vast majority of the 356 binding sites are clustered closely together with the remaining 20 sites scattered far away from the others, which is sensible based on the MDS plot of the full data in Fig 7. Since these results suggest problems with data quality, we did not include these outlying observations in the construction of our models and instead built the models using the remaining 906 binding sites. We will, however, present classification results both with and without the removed observations.

**Fig 8.**
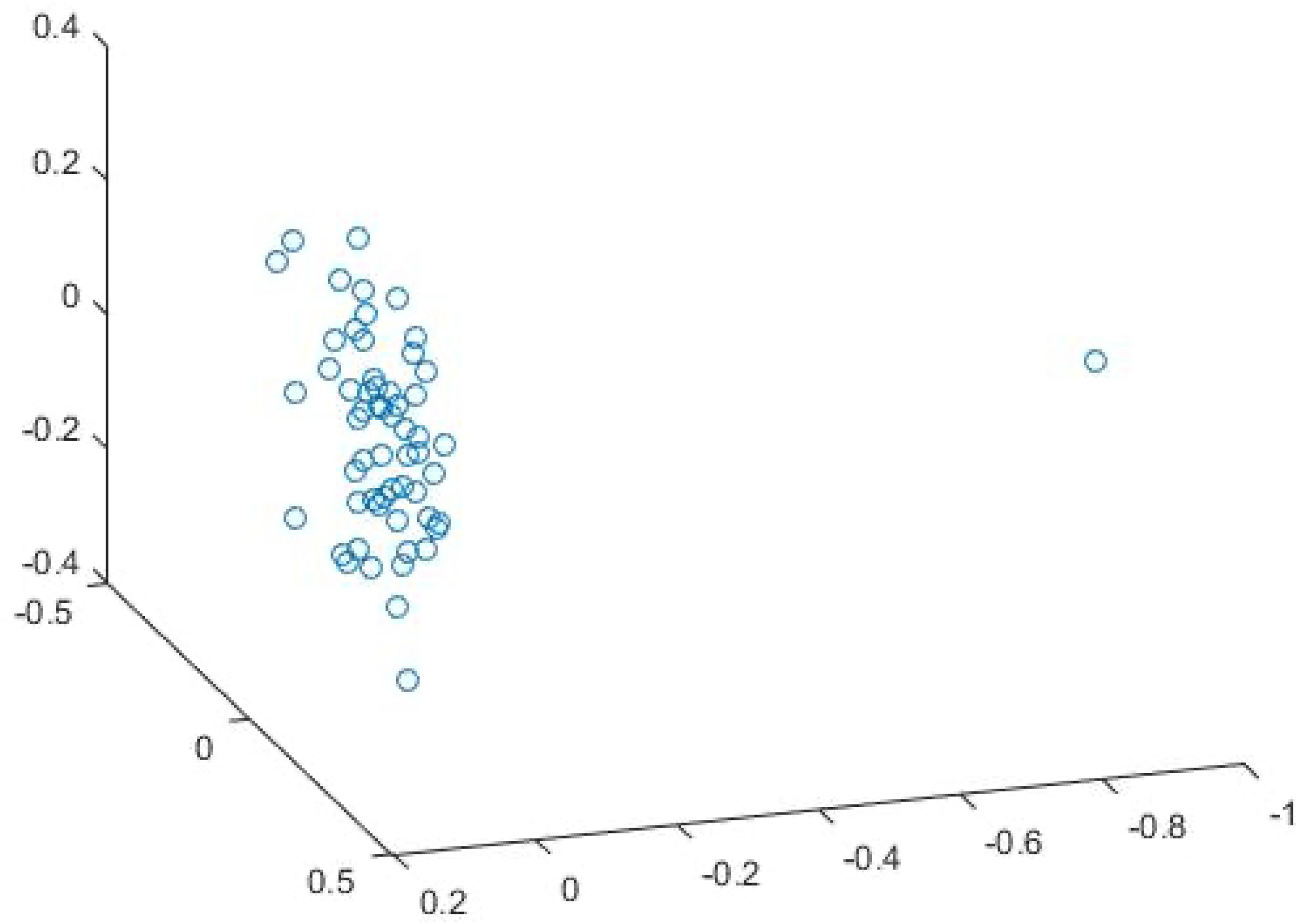
MDS plot in 3 dimensions for the FMN group of the extended Kahraman dataset before cleaning the data.

**Fig 9.**
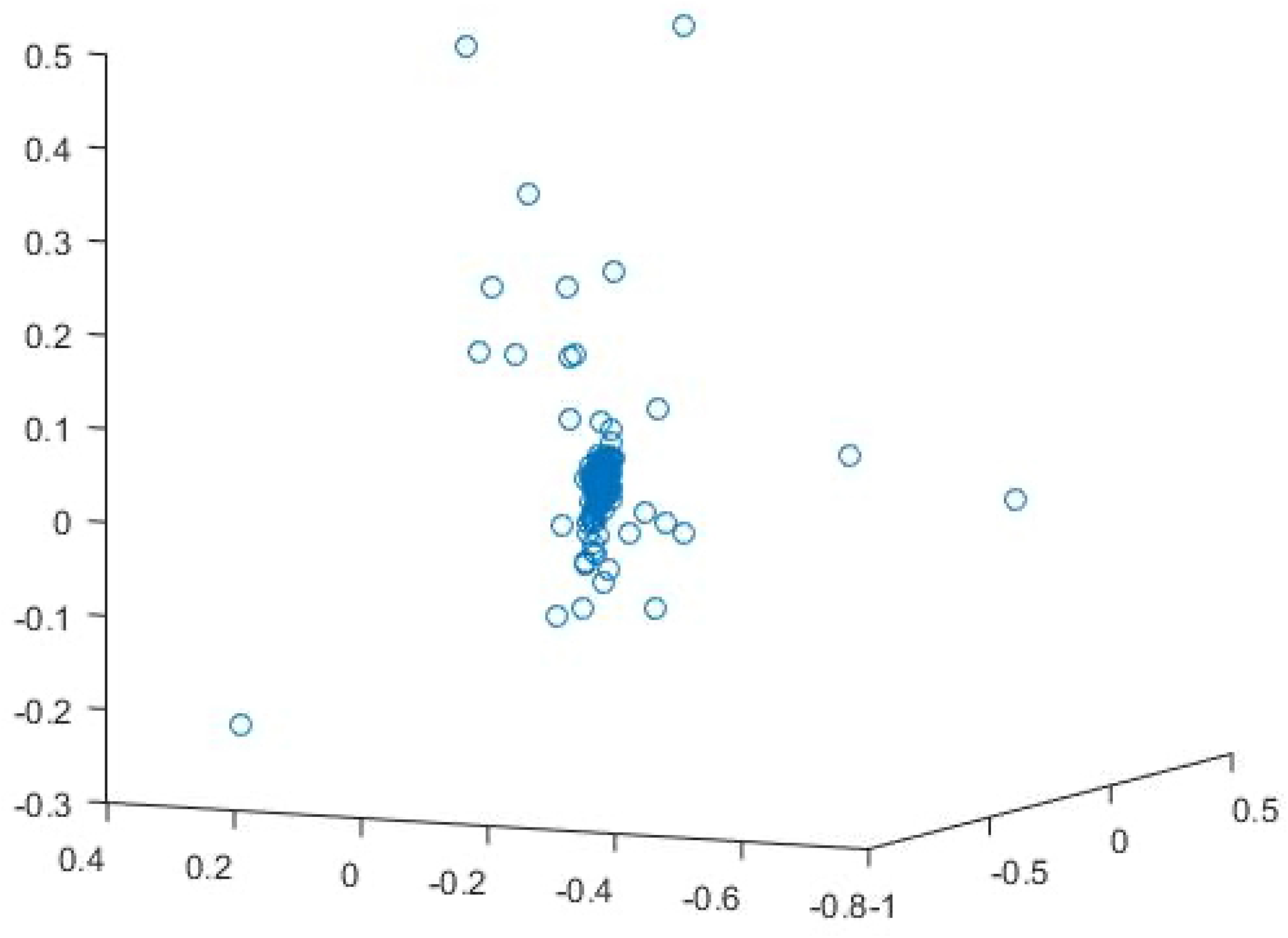
MDS plot in 3 dimensions for the PO4 group of the extended Kahraman dataset before cleaning the data.

While MDS plots for most of the remaining ligand groups revealed no issues with the data, we did notice an interesting pattern in the HEM group, as shown in Fig 10. Similar to the PO4 group, we see two distinct groups of observations, most of which are clustered along the right side of this plot with the remaining ones forming a long tail with variability almost entirely in just two of the three dimensions of this plot.

**Fig 10.**
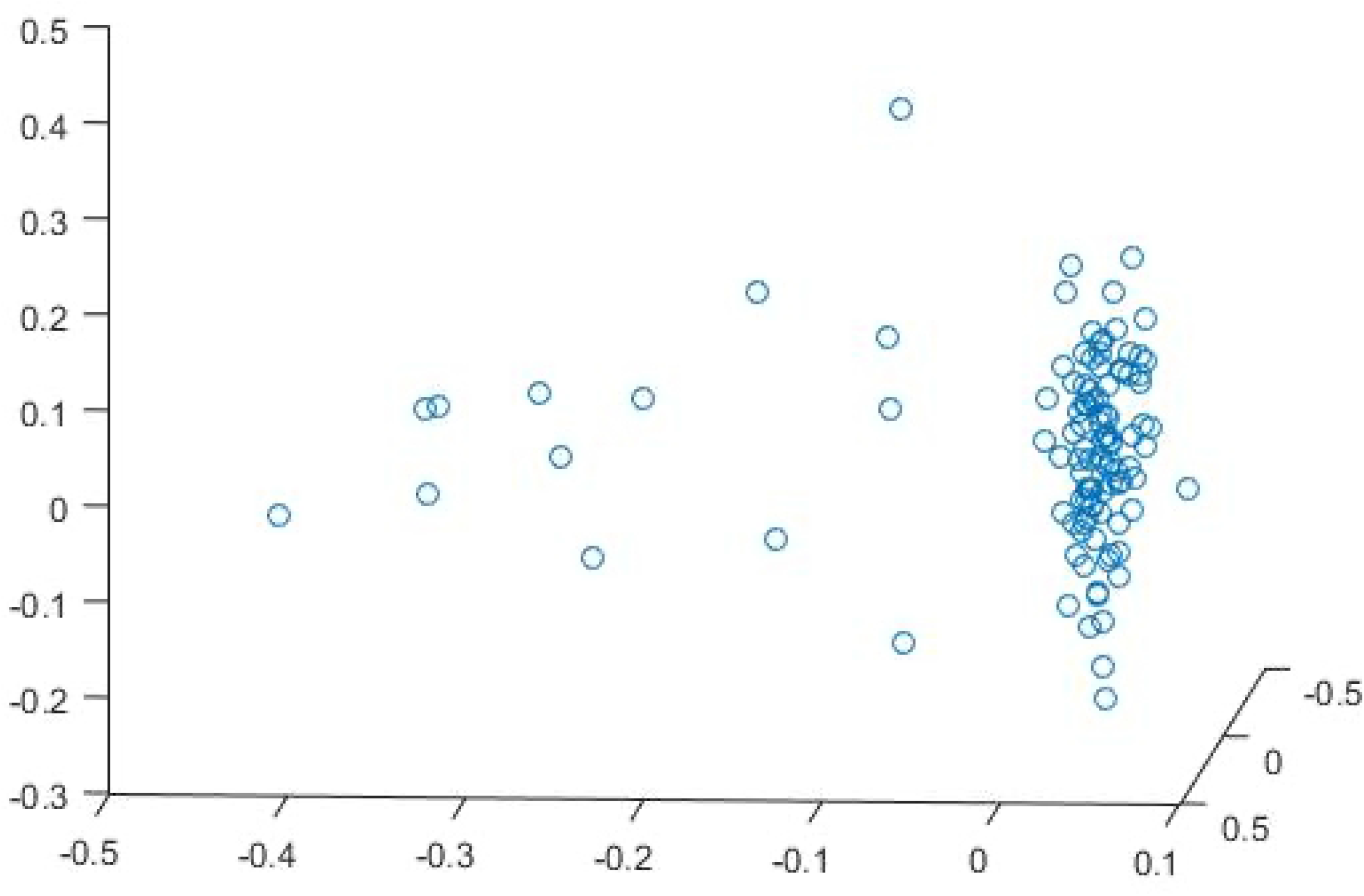
MDS plot in 3 dimensions for the entire HEM group of the extended Kahraman dataset.

To address this clustering problem, we broke the HEM group into two subsets, which we refer to as HEM-I and HEM-II. The MDS plots in 3 dimensions for these groups, shown in Fig 11 and 12, respectively, show that their subgrouping will have means and covariance matrices that more accurately capture the distributions of these binding sites. Please note that we could not use the same approach with the PO4 group since the space spanned by the cleaned observations is so large that Mahalanobis distances to the mean of those observations would be extremely small compared to the distances to the means of the other groups. Here, though, we see clear and distinct patterns that allow us to partition the HEM group for constructing the classification models. For evaluating the classification results, though, we still treat HEM as one entire group.

**Fig 11.**
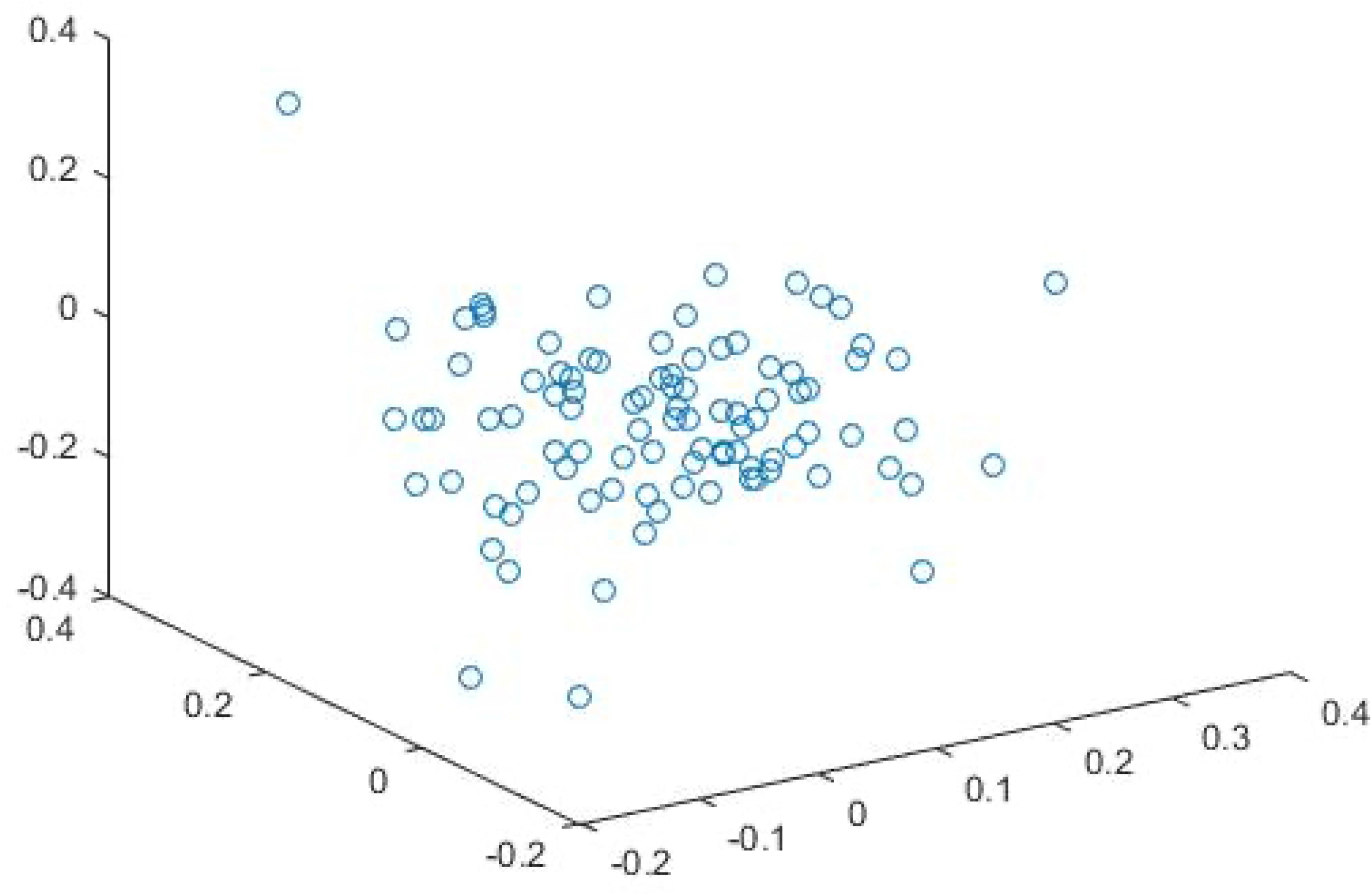
MDS plot for ligand HEM-I of the extended Kahraman dataset

**Fig 12.**
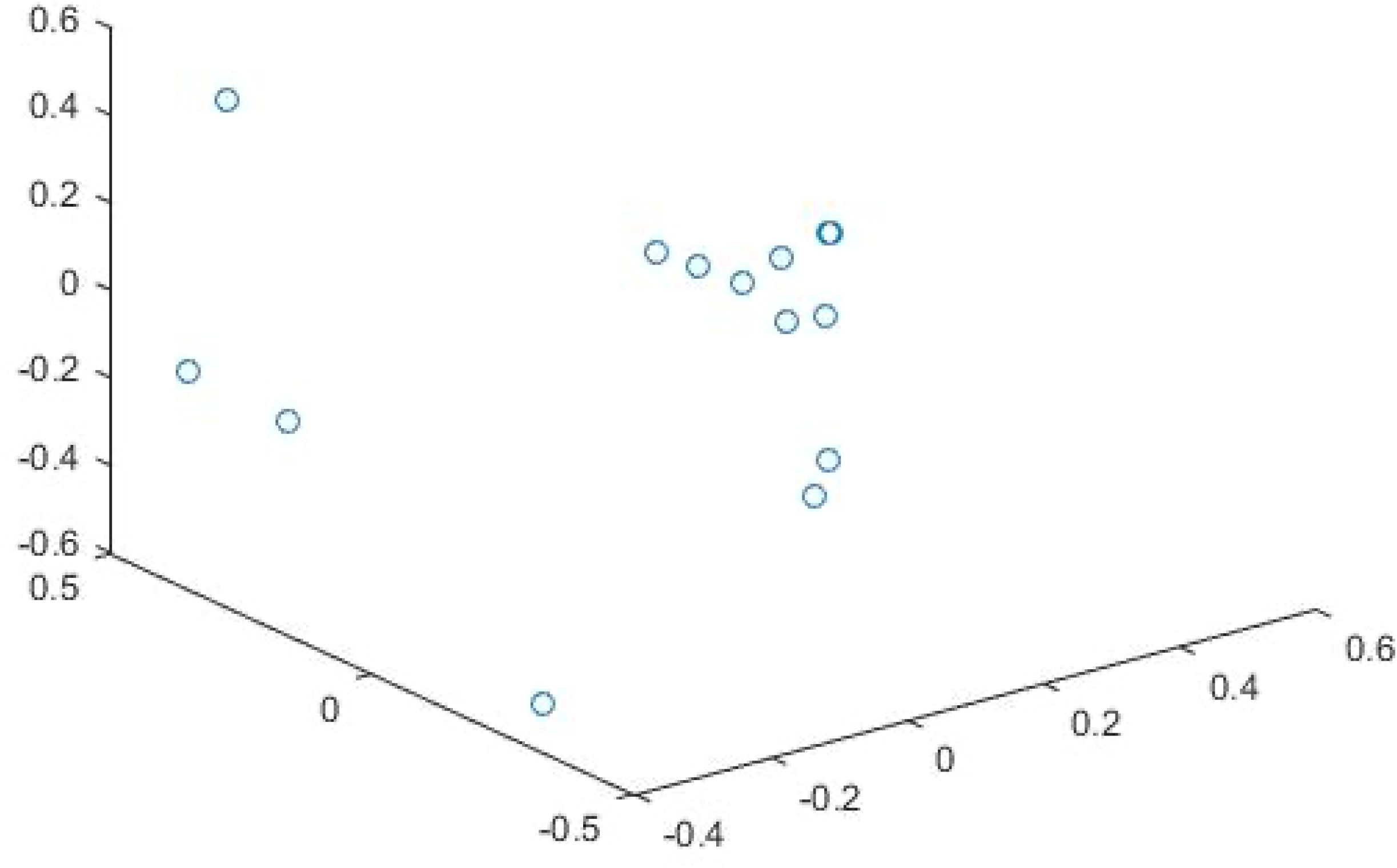
MDS plot for ligand HEM-II of the extended Kahraman dataset

An MDS plot in 2 dimensions of the entire cleaned dataset is shown in Fig 13. The patterns shown in this plot closely resemble those shown in Fig 5, though one axis is flipped, as commonly happens with MDS. While it was not our intention to obtain a similar distribution of observations for this dataset, this does provide support that the cleaning of the data was indeed necessary. From this, it is also clear that we did not over-clean the PO4 group since we still see far more variation in the group for this dataset than we did for the smaller set and that we simply removed observations with clear quality issues. Additionally, the clear separation between the two HEM subgroups is even clearer when viewed in combination with the rest of the data.

**Fig 13.**
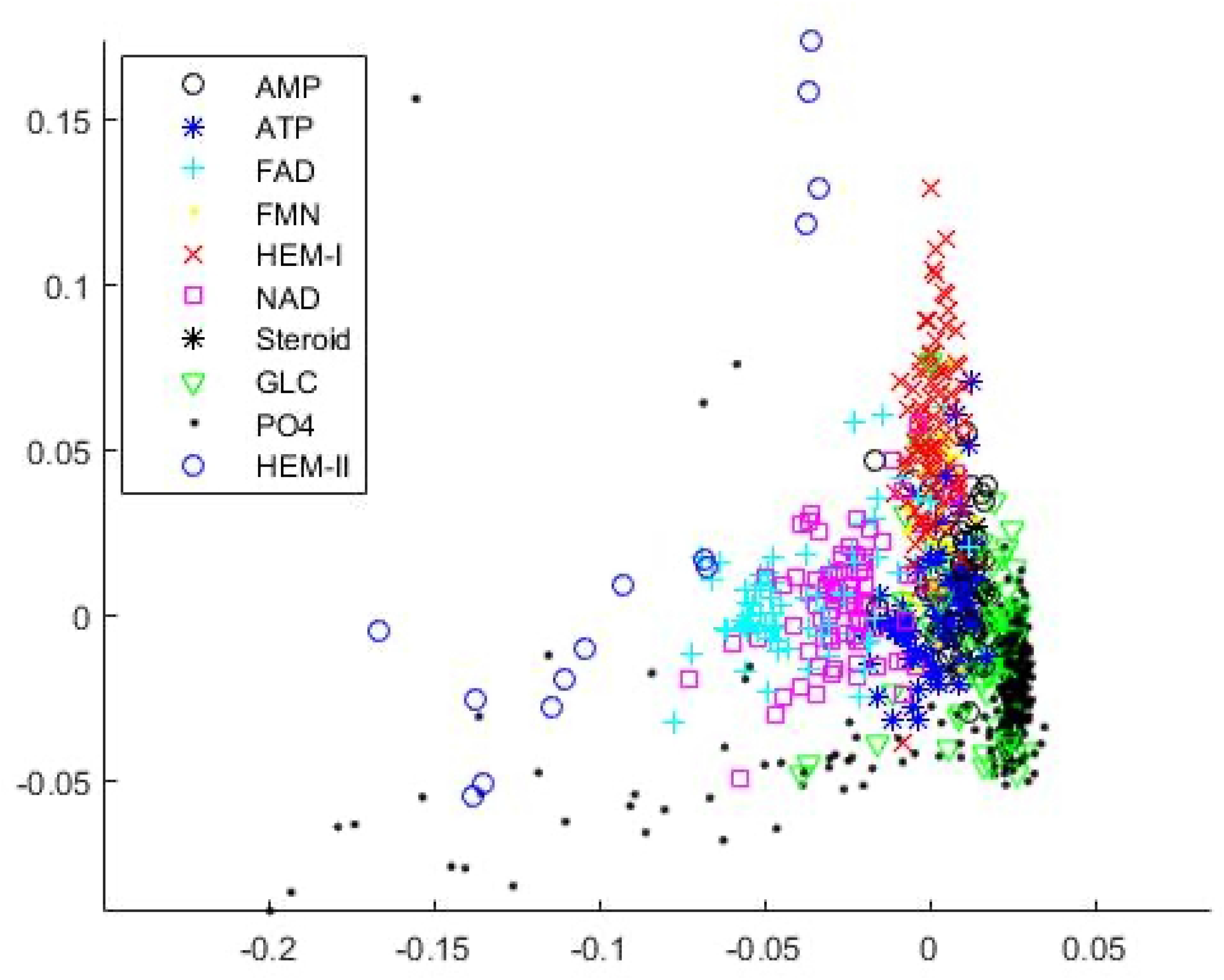
MDS plot in 2 dimensions of the entire cleaned extended Kahraman dataset.

A summary of the results of the classification study for this dataset is shown in Table 5. Results are not reported for the TIPSA variants because of the extreme computational cost of that method for a dataset of this size. Because some proteins from the original extended Kahraman set were not still available on PDB and we performed light data cleaning, we cannot perfectly compare the classification results of CDPA with those of [14]. Despite this, though, we still present their results to provide a rough baseline for performance and to highlight differences between the methodologies between their study and ours.

**Table 5.** Results for nearest neighbor classification for the extended Kahraman (5.3 Å) dataset.

| Method | Classification Error |
| --- | --- |
| *Sup - CK<sub>L</sub>* | 0.19 |
| **CDPA with nearest mean classification**<br>**(When trained and tested with 906 obs.)** | **0.32** |
| **CDPA with nearest mean classification**<br>**(When trained with 906 obs. and tested with 965 obs.)** | **0.3451** |
| **CDPA with polytomous logistic regression** | **0.25** |

Unlike for the Kahraman dataset, here, CDPA produces higher classification errors than Sup-CK does. The difference is especially clear for the nearest mean classifier, which performs worse than the logistic regression classifier. A primary reason for the difference in performance between Sup-CK and CDPA is that the former method utilizes pairwise comparisons between observations while ours uses model-based comparisons to means of each group. While an MDS plot for Sup-CK’s Gaussian convolution kernel encoding of the binding sites would likely differ at least somewhat from that of CDPA, the general principle would remain the same, so we will appeal Fig 13 to highlight differences between the methods. The PO4 observation with coordinates of roughly (−0.19, −0.09) is misclassified under our nearest mean classifier because it is on the extreme edge of the distribution of PO4 and, as such, has a high Mahalanobis distance to PO4’s mean. On the other hand, though, its five nearest observations are to other PO4 sites, so a k-nearest neighbor classifier would classify it correctly. This would also be true of many of the PO4 binding sites shown in this plot as well as the PO4 observations that were cleaned from the data. The use of the Mahalanobis distances, though, allows for correct classification of many other binding sites with CDPA, though, which k-nearest neighbor methods may fail for.

We now present a detailed analysis of the results for CDPA for both classification methods, beginning with the nearest mean classifier. The confusion matrix for this method is shown in Table 6. A number of the misclassifications are sensible due to the similarity of the binding ligands. For instance, AMP and ATP are quite similar structurally, show it is not surprising to see that many binding sites from the AMP and ATP groups are incorrectly classified as belonging to the other group. The performance and similarity of the ligands for FAD and NAD is likewise. While the structures of GLC and PO4 are not so similar, they are both by far the smallest ligands in this data set, so it is not surprising that sites from these two groups may be mistaken for each other.

**Table 6.** Classification of binding sites for extended Kahraman dataset using nearest mean classification

|  | AMP | ATP | FAD | FMN | HEM | NAD | Steroid | GLC | PO4 |
| --- | --- | --- | --- | --- | --- | --- | --- | --- | --- |
| AMP(61) | **33** | 13 | 1 | 0 | 2 | 1 | 0 | 8 | 3 |
| ATP(77) | 11 | **44** | 0 | 5 | 2 | 5 | 0 | 2 | 8 |
| FAD(78) | 0 | 5 | **58** | 0 | 2 | 12 | 0 | 0 | 1 |
| FMN(56) | 0 | 19 | 0 | **23** | 8 | 1 | 0 | 4 | 1 |
| HEM(109) | 2 | 5 | 1 | 5 | **93** | 1 | 0 | 0 | 2 |
| NAD(88) | 0 | 3 | 23 | 2 | 0 | **59** | 0 | 0 | 1 |
| Steroid(6) | 1 | 0 | 0 | 0 | 0 | 0 | **5** | 0 | 0 |
| GLC(75) | 1 | 7 | 0 | 0 | 2 | 1 | 0 | **44** | 20 |
| PO4(356) | 2 | 3 | 3 | 0 | 7 | 2 | 0 | 81 | **258** |

Even though we didn’t see this behavior from the Kahraman data set, it is easy to see why these groups were misclassified for each other here from Fig 13; the two groups are close neighbors to each other, often overlapping, especially in the long tail of the PO4 group that extends to the left side of the plot.

We now consider the logistic regression classification scheme, for which we used the ligand group AMP as the reference category for calculating log-odds ratios with respect to. We then calculated predicted probabilities that each observations belong to the ligands groups based on these. As shown in Table 5, the classification error for this method is closer to that of Sup-CK than to CDPA with the nearest mean classifier, which provides support for the idea that it is important to utilize Mahalanobis distances to the means of all groups rather than just consider which mean is closest. Table 7 shows the estimated coefficients for every parameter in the poloytomous logistic regression model. Since the predictor variables are distances, negative coefficients decrease the log-odds that a binding site belongs to that group compared to AMP for every increase in the distance of an observation to that group. As expected, then, the bold elements along the diagonal of the table are negative. As a further example for interpreting the coefficients, consider the HEM-II group; these coefficients explain that if the binding site is closer to AMP, ATP, FAD, GLC, and HEM-II, but further away from FMN, HEM-I, NAD, Steroid, and PO4, it is more likely to be coming from HEM-II than AMP.

**Table 7.** Coefficients of the polytomous logistic regression model for the extended Kahraman dataset.

|  | Predictors |  |  |  |  |  |  |  |  |  |  |
| --- | --- | --- | --- | --- | --- | --- | --- | --- | --- | --- | --- |
| Response | Intercept | AMP | ATP | FAD | FMN | HEM-I | NAD | Steroid | GLC | PO4 | HEM-II |
| ATP | -1.43 | 2.59 | **-2.68** | 0.73 | 0.27 | -0.25 | -0.44 | -0.03 | -0.33 | 0.29 | 0.06 |
| FAD | 0.54 | 3.75 | 1.33 | **-3.56** | -0.74 | 0.11 | 0.65 | -0.16 | -0.27 | -0.81 | 0.28 |
| FMN | 1.51 | 2.70 | -0.22 | 0.87 | **-2.16** | -0.07 | -1.21 | -0.06 | -0.68 | 0.29 | 0.15 |
| HEM-I | 2.08 | 1.78 | 0.52 | 0.95 | 1.10 | **-3.23** | -1.15 | -0.04 | -0.07 | -0.18 | 0.05 |
| NAD | 2.57 | 2.69 | -0.76 | -0.93 | 0.11 | 0.48 | **-2.47** | -0.03 | -0.29 | 0.38 | -0.16 |
| Steroid | 83.01 | 81.53 | -156.18 | -779.05 | -152.96 | 65.35 | 997.54 | **-56.99** | 187.35 | -171.69 | -60.03 |
| GLC | -2.63 | 2.99 | -1.02 | 0.11 | 0.41 | -0.11 | 0.84 | -0.04 | **-2.42** | 0.26 | -0.27 |
| PO4 | -6.72 | 3.32 | 0.06 | -2.17 | 0.62 | -0.62 | 2.25 | -0.08 | -0.89 | **-1.62** | 0.56 |
| HEM-II | 30.08 | -8.85 | -105.65 | -1.83 | 35.46 | 16.22 | 17.25 | 3.72 | -33.35 | 51.78 | **-31.95** |

The confusion matrix for the logistic regression classifier is shown in Table 8. The overall pattern is very similar to what we saw for the nearest mean classifier. However, the biggest difference between them is in the trade-off between the PO4 and GLC groups. While a higher percentage of GLC sites are now misclassified as being from the PO4 group, nearly all of the PO4 sites are now correctly classified.

**Table 8.** Classification of binding sites for extended Kahraman dataset using the logistic regression classifier

|  | AMP | ATP | FAD | FMN | HEM | NAD | Steroid | GLC | PO4 |
| --- | --- | --- | --- | --- | --- | --- | --- | --- | --- |
| AMP(61) | **35** | 8 | 1 | 6 | 0 | 1 | 1 | 5 | 4 |
| ATP(77) | 12 | **45** | 1 | 8 | 1 | 5 | 0 | 4 | 1 |
| FAD(78) | 1 | 2 | **54** | 2 | 3 | 14 | 0 | 0 | 2 |
| FMN(56) | 2 | 11 | 0 | **33** | 10 | 0 | 0 | 0 | 0 |
| HEM(109) | 2 | 2 | 1 | 5 | **98** | 0 | 0 | 0 | 1 |
| NAD(88) | 0 | 3 | 14 | 1 | 1 | **69** | 0 | 0 | 0 |
| Steroid(6) | 0 | 0 | 0 | 0 | 0 | 0 | **6** | 0 | 0 |
| GLC(75) | 3 | 4 | 0 | 2 | 2 | 1 | 0 | **17** | 46 |
| PO4(356) | 4 | 2 | 2 | 0 | 1 | 0 | 0 | 7 | **340** |

While our analysis of the nearest mean classifier stopped with understanding the confusion matrix, we can go further with the logistic regression classifier because it produces predicted probabilities that all observations belong to each class. The predicted probability for the assigned group is most informative because it provides a measure of certainty for each prediction. That is, if a predicted probability is high, it provides a higher degree of certainty in the prediction based on the model. As such, we should hope that the predicted probabilities for the correctly classified sites tend to be higher than those for the incorrectly classified sites. Fortunately, this is what we observe in comparative boxplots of the distributions of the predicted probabilities in Fig 14. 50% of the correctly classified sites have predicted probabilities over 0.9 and just 25% of these probabilities are below 0.7. This is in sharp contrast with the predicted probabilities for the misclassified sites, of which only 25% are above 0.7.

**Fig 14.**
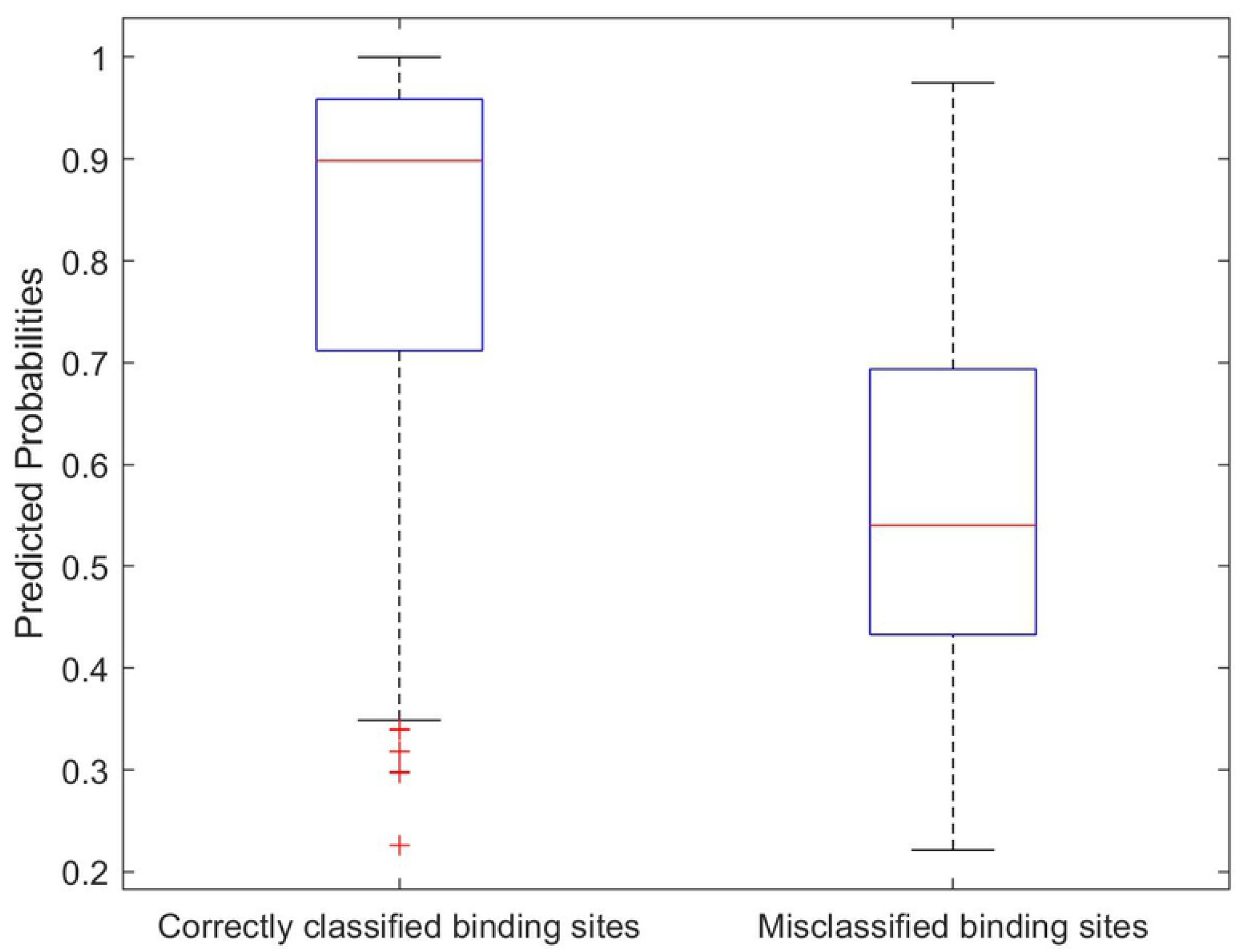
Correctly classified and misclassified binding sites vs their predicted probabilities. A comparative boxplots of the distributions of the predicted probabilities.

Fig 15 and 16 display the distributions of the predicted probabilities for the correctly and incorrectly classified binding sites, respectively, for each ligand group. Binding sites from the ligand groups AMP, FAD, FMN, HEM, NAD, Steroid, and PO4 tend to have higher predicted probabilities for correct classifications than for the incorrect classifications. These two figures also highlight that most of the misclassified sites from the GLC group were misclassified with high predicted probabilities. This result is not surprising since Table 8 shows that most of the GLC sites were misclassified as PO4 sites and Fig 13 shows that the PO4 and GLC groups overlap.

**Fig 15.**
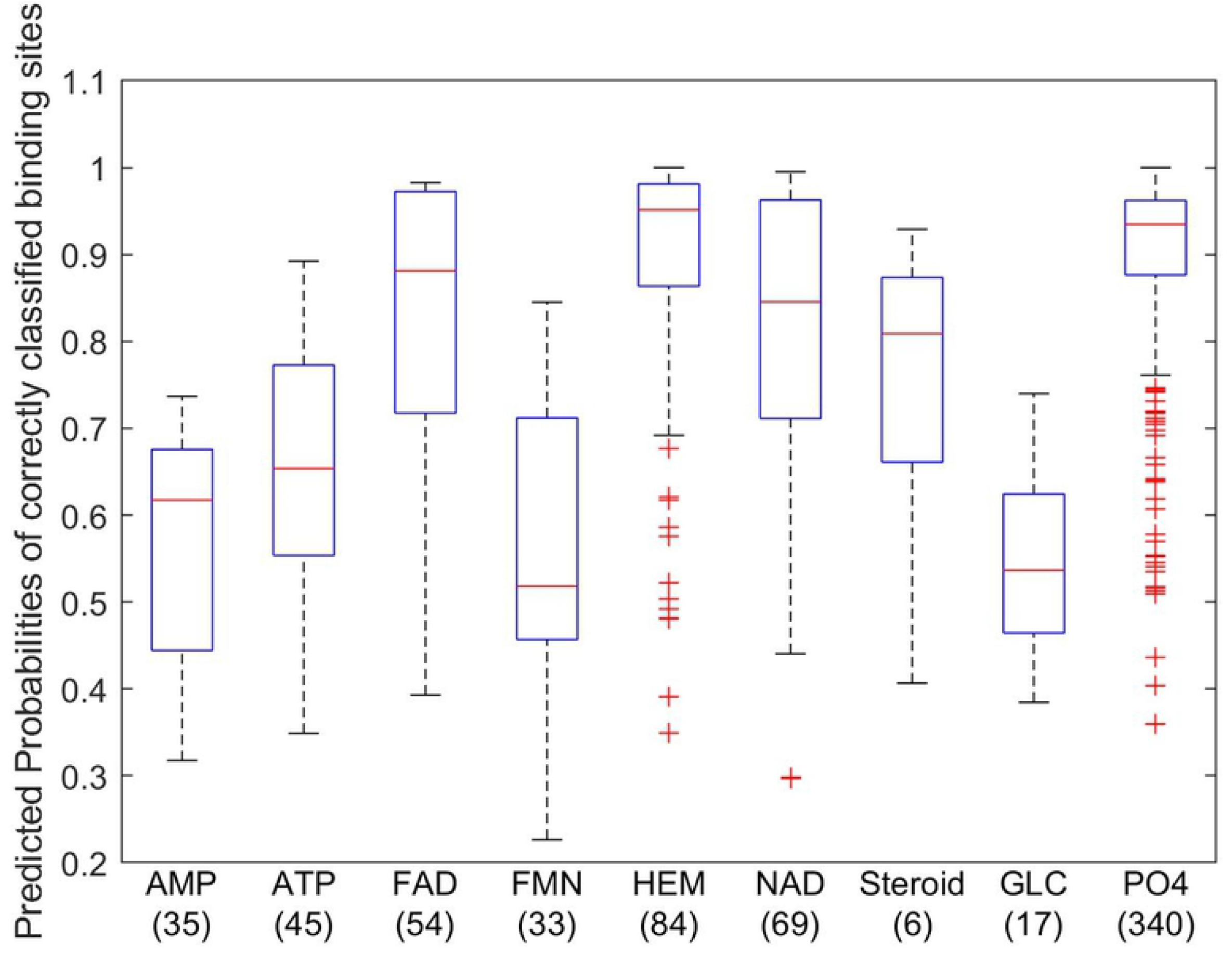
Predicted Probabilities of correctly classified binding sites of each ligand groups. A comparative boxplots that display the distributions of the predicted probabilities for the correctly classified binding sites.

**Fig 16.**
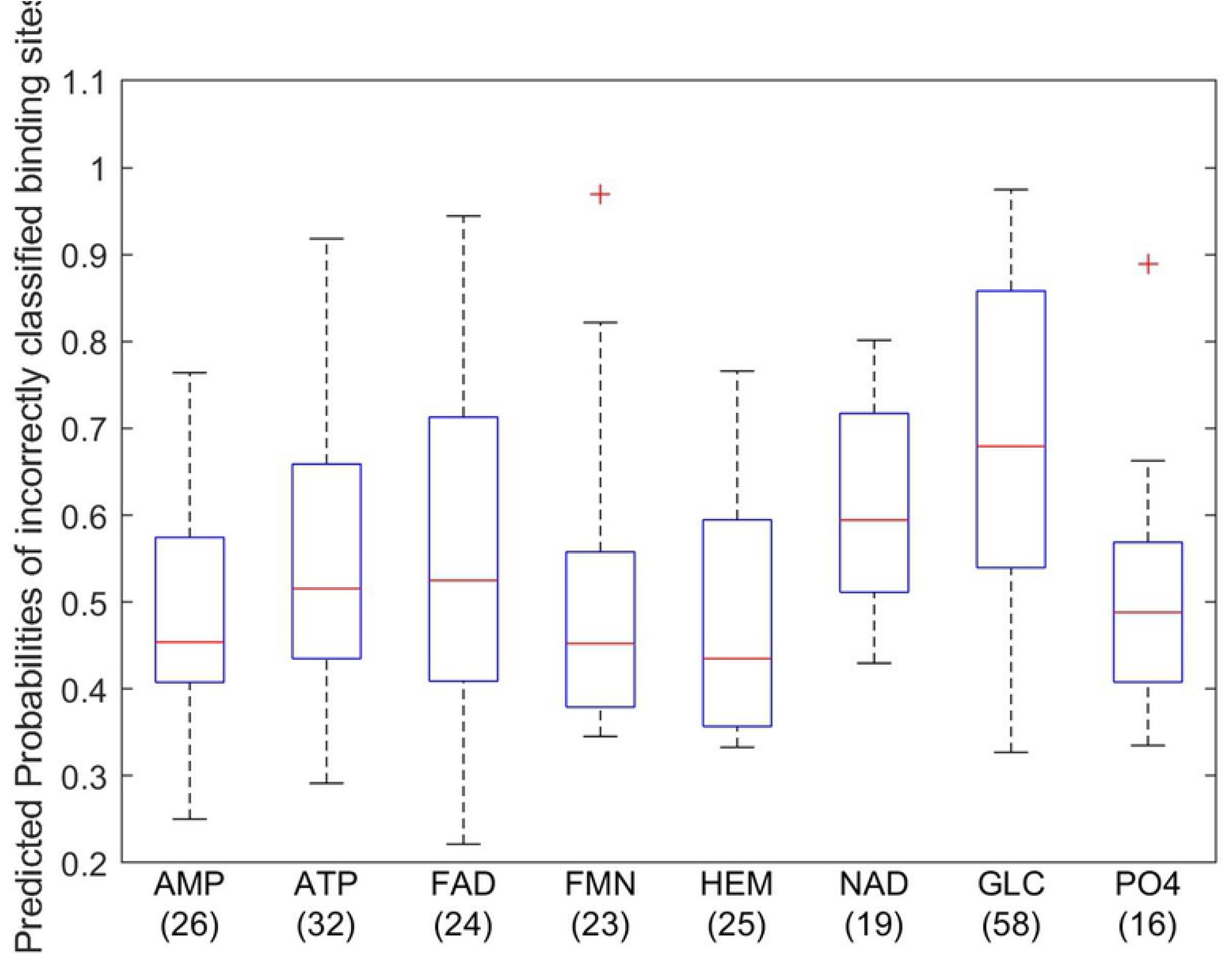
Predicted Probabilities of incorrectly classified binding sites of each ligand groups. A comparative boxplots that display the distributions of the predicted probabilities for the incorrectly classified binding sites.

To perform validation for the CDPA classification procedures, we once again added varying amounts of zero-mean Gaussian noise to the coordinates of the binding sites prior to encoding the data as covariance matrices. Fig 17 visually compares the classification errors for both the nearest mean and logistic regression classifiers as a function of the level of noise added to the data. Compared to the results for the Kahraman set, these procedure are both far less sensitive to even small amounts of noise since the original dataset is much larger here. The average classification error for the nearest mean classifier increase roughly linearly with the noise level, but for the logistic regression classifier, the average classification error first increases quite slowly for noise levels up to roughly 0.3 and then increases roughly linearly for noise levels above that.

**Fig 17.**
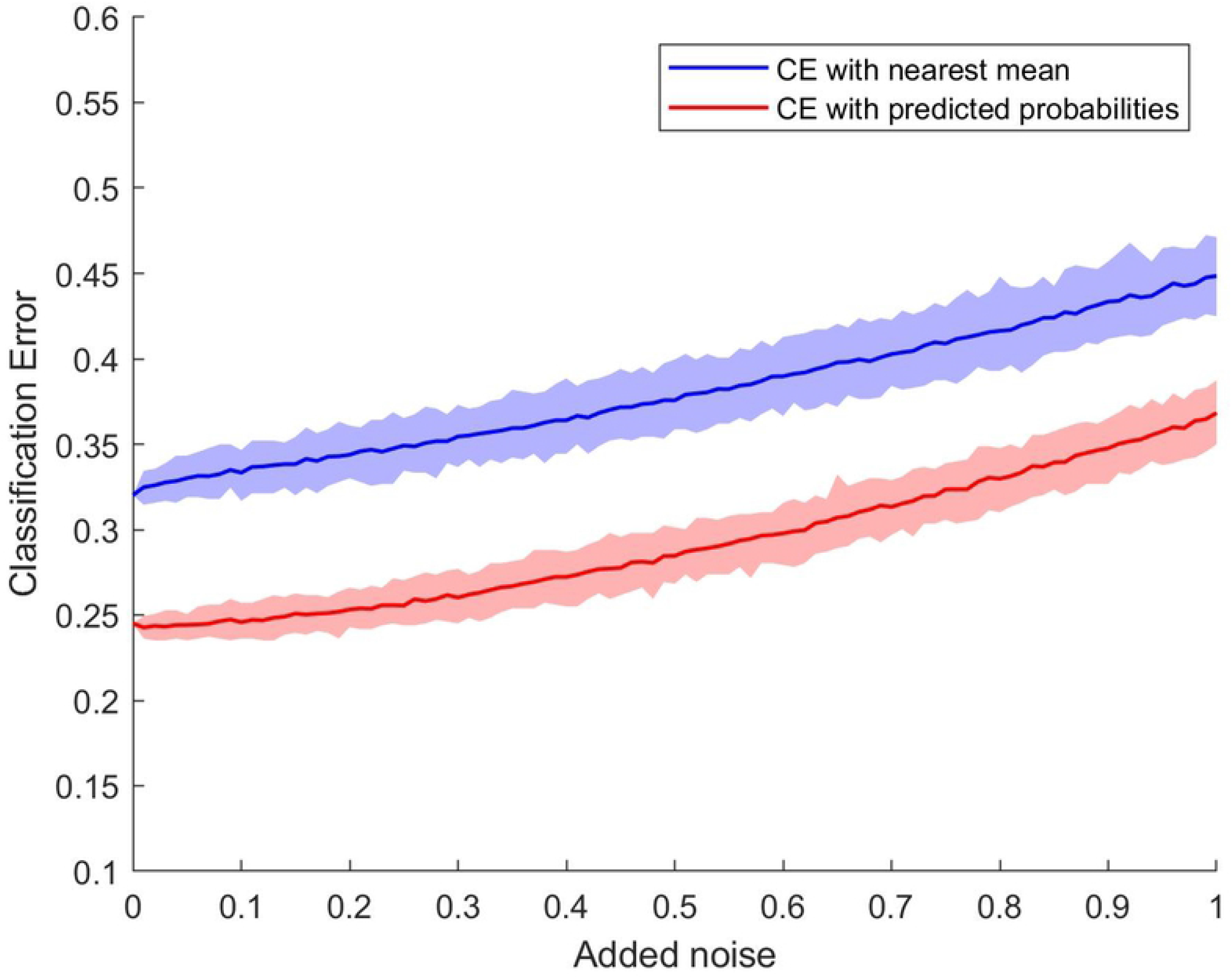
Average classification errors (dark lines) and their 95% confidence bands for the simulated validation sets. classification errors for both the nearest mean and logistic regression classifiers as a function of the level of noise added to the data.

These results further confirm the superiority of the logistic regression classifier to the nearest man classifier since the simulated classification error for the former method with noise levels near 1 is comparable to the classification error of the nearest mean method even for extremely low levels of noise. Furthermore, the confidence bands for the logistic regression classifier are considerably narrow at all levels of noise.

### Comparison of computational costs

The model-based approach used in this research is very fast compared to the alignment-based approaches used by [15] and [14]. For instance, in [15], to perform pairwise alignment for the Kahraman dataset, it took between 2 to 6 seconds per alignment. Since there are 100 binding sites in this dataset, there will be 4950 pairs, requiring a minimum of 9900 seconds to compare all of the pairs. For [14], the algorithm running time per pockets pair varied between 0.2 and 1.3 seconds. This means that, at a minimum, it will take 990 seconds to compare all the pairs using that methodology. With CDPA, to calculate the similarity measure for all binding sites and classify them to their ligand group, it took only 2.4 seconds for the Kahraman dataset. The extreme difference in computational costs is due to both CDPA being alignment-free and our classifiers not being restricted to pairwise comparisons. The difference in computation times for the extended Kahraman dataset would be even more striking due to the considerably larger sample size requiring two orders of magnitude more pairwise comparisons.

## Conclusion

In this study, we developed a novel representation called CDPA for encoding the structural information from a protein binding site as a 3 × 3 covariance matrix. This representation allowed us to develop nonparametric probability models for groups of sites that all bind to the same ligand using the Mahalanobis distance for the covariance matrix for each binding site to the mean of the binding group. We then showed that these distributions of the distances are useful for classifying the sites by binding ligand. CDPA with the nearest mean classifier outperformed others methods for the Kahraman dataset. While it is improper to compare directly to the results of [14] for the extended Kahraman dataset since some of the set’s proteins are no longer listed in the PDB and we had to perform some light data cleaning, it is clear that CDPA with the logistic regression classifier still performed comparably to [14] for the dataset. At the very least, this suggests the CDPA is able to discriminate between the ligand groups. At the same time, CDPA is orders of magnitude faster than the alignment-based methods since it is invariant to coordinate changes for the atoms and does not rely on pairwise comparisons, but instead on comparisons to the mean covariance matrix for each ligand group.

The differences between pairwise and model-based comparisons also extend to classification performance. For instance, sup-CK with the nearest neighbor classifier performs well for the extended Kahraman dataset, despite some data quality issues, because the outlying observations only influence classification performance for each other since no models for the groups are constructed. This is in sharp contrast to CDPA models, where the outlying observations also impact the construction of the models when not cleaning the data first. On the other hand, though, nearest neighbor methods are overly sensitive to noise and cannot take variation within and across binding site groups into considerations, whereas our model-based approaches do so explicitly.

To explore how our CDPA classifiers performed with different data, we simulated testing sets by adding increasing amounts of Gaussian noise to the coordinates of the atoms in the binding sites. Fig 6 and 17 showed the average classification error per replication with confidence bands as a function of the amount of noise for the two datasets we used to form the models. While the classification error rate was lower for the original Kahraman dataset than for the extended Kahraman dataset, the classification performance degraded more quickly as noise was added to the data. Furthermore, in Fig 18, we can see that the lengths of the confidence intervals for the extended Kahraman data are much narrower than those of the Kahraman set. As a result, we can be more confident that the CDPA procedures are, in fact, more stable with more observations, even though the performance for the larger dataset is not quite as good as it is for the smaller set. This provide further support for the ability of CDPA to capture differences between groups of binding sites.

**Fig 18.**
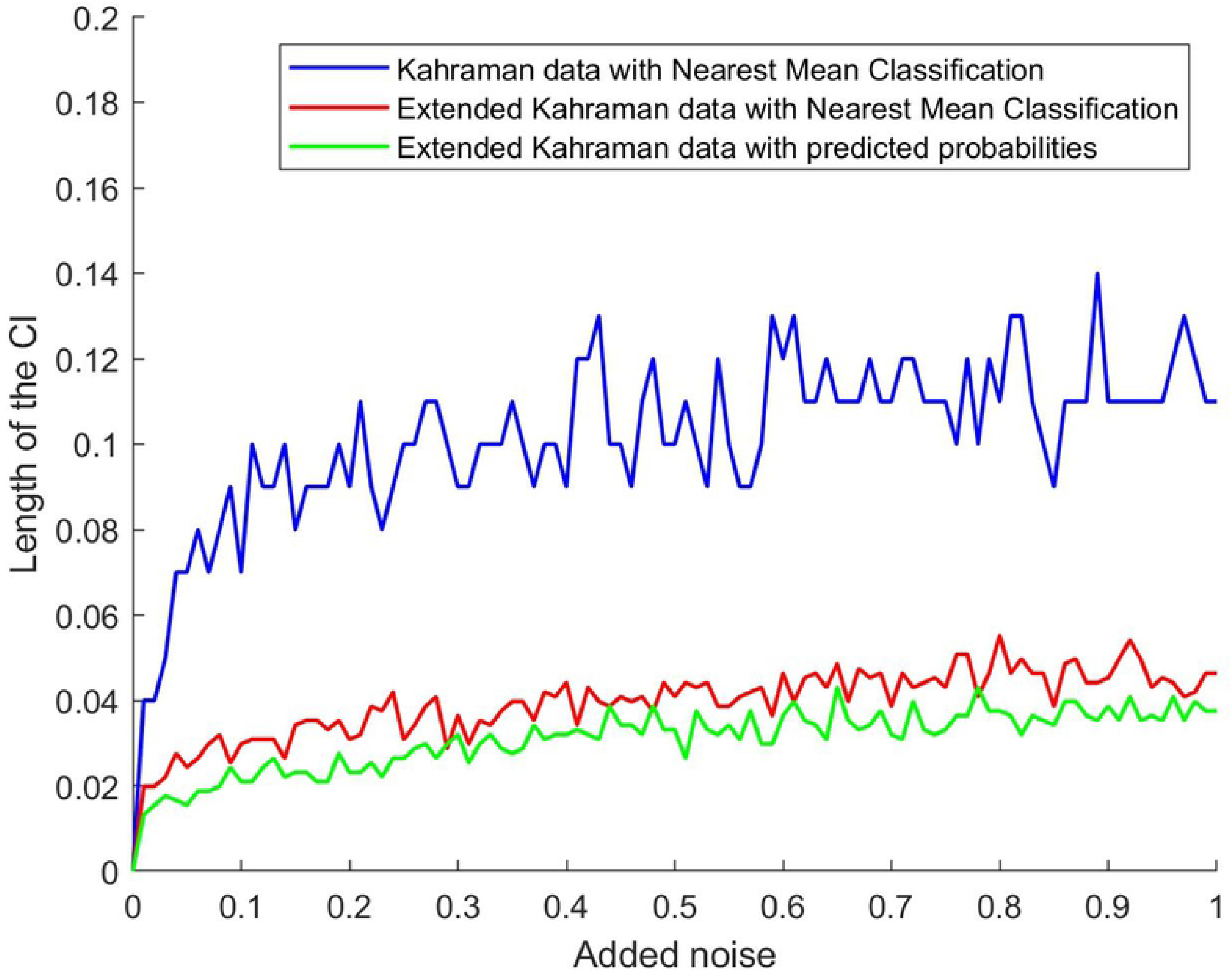
Lengths of the confidence intervals for the Kahraman and the extended Kahraman datasets as a function of noise level.

However, there is plenty of work that remains to be done to build upon this initial study on CDPA. As we mentioned above, the CDPA models appear to work better for those binding sites having similar covariance structures to the average of their group, but pairwise comparisons appear to be more advantageous for outlying sites. This suggests that future approaches could seek to leverage both types of classification methods: model-based for binding sites with relatively low distances to at least one group and pairwise comparisons for those observations that differ considerably to the means of the groups. Furthermore, we wish to consider additional classification procedures, such as classification trees, parametric probability models that would permit Bayesian procedures to be used, and ensemble methods that could combine many or all of the approaches.

Ideally, these methods would permit us to also utilize alternative and/or additional dissimilarity measures to what we considered here. For instance, in this paper, we utilized the fact that the space of 3 × 3 covariance matrices is a submanifold of the space of 3 × 3 symmetric matrices to utilize the Euclidean distance between observations. However, there are many additional metrics that can be placed on the space of 3 × 3 covariance matrices that would define other distances between observations. The other distances may be able to further improve the performance of CDPA by more explicitly taking the geometry of the sample space into consideration. Additionally, we seek to enhance CDPA by bringing other characteristics of the binding sites, especially chemical attributes, into the analysis, as well, since that information is not explicitly taken into account by CDPA.

